# TAT-Cx43_266-283_ impairs metabolic plasticity in glioma stem cells in vitro and in vivo

**DOI:** 10.1101/858563

**Authors:** Sara G. Pelaz, Myriam Jaraíz-Rodríguez, Rocío Talaverón, Laura García-Vicente, Marta Gómez de Cedrón, María Tabernero, Ana Ramírez de Molina, Concepción Lillo, José M. Medina, Arantxa Tabernero

## Abstract

Glioblastoma is the most aggressive primary brain cancer, with a median survival of 1 to 2 years^1^. These tumours contain glioma stem cells (GSCs), which are highly tumorigenic, resistant to conventional therapies^2, 3^, and exhibit metabolic plasticity to adapt to challenging environments^4, 5^. GSCs can be specifically targeted by a short cell-penetrating peptide based on connexin43 (Cx43) (TAT-Cx43_266-283_) that reduces tumour growth and increases survival in preclinical models^6^ via c-Src inhibition^7^. Because several reports revealed poor clinical efficacy of various antitumoral drugs due to metabolic rewiring in cancer cells^8–10^, we investigated the effect of TAT-Cx43_266-283_ on GSC metabolism and metabolic plasticity. Here we show that TAT-Cx43_266-283_ decreases GSC glucose uptake and oxidative phosphorylation without a compensatory increase in glycolysis, with no effect on neuron or astrocyte metabolism. GSC changes were mediated by decreased hexokinase (HK) activity and aberrant mitochondrial localization, ultrastructure and function. Moreover, TAT-Cx43_266-283_ reduced GSC growth and survival under different nutrient availability conditions by impairing the metabolic plasticity needed to exploit glucose as an energy source in the absence of other nutrients. Finally, GSCs intracranially implanted into mice together with TAT-Cx43_266-283_ showed decreased levels of important targets for cancer therapy, such as HK-2^11, 12^ and glucose transporter 3 (GLUT-3)^13^, evidencing the reduced ability of treated GSCs to survive in challenging environments. Our results confirm the value of TAT-Cx43_266-283_ for glioma therapy alone or in combination with therapies whose resistance relies on metabolic adaptation. More importantly, these results allow us to conclude that the advantageous metabolic plasticity of GSCs is a targetable vulnerability in malignant gliomas.

## Introduction

Glioblastoma (World Health Organization grade IV glioma) is the most common and aggressive primary brain cancer, with a median survival of 16 months^1^. A subset of cells within these tumours, termed glioma stem cells (GSCs), display highly tumorigenic capacity and resistance to conventional therapies, and are therefore considered responsible for recurrence^2, 3^. The GSC phenotype is dynamic and can be acquired by non-GSCs upon challenging conditions, suggesting that inherent cancer cell plasticity is a relevant target for treatment^13, 14^. These challenging conditions, which include hypoxia, an acidic environment or metabolic stress, promote the enrichment of the highly tumorigenic GSC population^13, 15, 16^. In this regard, an important feature of cancer stem cells, and particularly of GSCs, is their metabolic plasticity, which allows them to survive nutrient deprivation by conveniently shifting between different metabolic pathways used in energy production and catabolism^4, 5, 17–19^. In particular, glucose metabolism plays a key role in glioma, as glucose restriction promotes GSC enrichment through a two-fold mechanism: direct selection of GSCs, because they have the metabolic phenotype required to survive in this environment, and adaptation of some non-GSCs through the acquisition of GSC metabolic features^13^. One of these GSC features is elevated expression of the high affinity glucose transporter, GLUT-3, which is crucial for tumorigenicity of GSCs, and consequently, for progression and malignancy of human gliomas^13^.

Characteristically, glioma cells, including GSCs, express low levels of connexin43 (Cx43), as opposed to astrocytes, one of their healthy counterparts^7, 20, 21^. In addition to forming intercellular gap junctions, Cx43 has gathered plenty of attention in recent years for its C-terminal domain^22^, which interacts with a plethora of molecules and acts as an intracellular signaling hub^23^. This is the case of the proto-oncoprotein c-Src, which is recruited by Cx43 together with its inhibitors CSK and PTEN^24^, causing the inhibition of c-Src and its oncogenic downstream pathways^25^. Based on this inhibitory mechanism, we designed a cell-penetrating peptide, TAT-Cx43_266-283_, containing the region of Cx43 required for the inhibition of c-Src^7^. Importantly, TAT-Cx43_266-283_ exerts potent antitumor effects on glioma cells^26^, including the reversion of GSC phenotype^7^ in vitro and in vivo without affecting healthy brain cells^6^.

The relationship between the oncogenic activity of c-Src and glucose metabolism was first described in 1985^27^. Since then, many studies have delved into the underlying mechanisms. c-Src can modulate glucose metabolism either indirectly through master transcription factors, such as c-myc^28^ or hypoxia inducible factor 1α (HIF-1α)^29, 30^, or directly through the regulation of the activity of key glucose metabolism proteins, such as HK-2^12^, which traps glucose intracellularly for further metabolization, or glucose-6-phosphate-dehydrogenase (G6PD)^31^, the key and rate-limiting enzyme in the pentose phosphate pathway.

In addition to a conferred advantage to survive in nutrient-deprived conditions, metabolic plasticity is responsible for resistance to certain cancer treatments, one of the most relevant hallmarks of GSCs^2, 3^. For example, glioma cells adapt to bevacizumab treatment – a common antiangiogenic therapy – by increasing glycolisis^8, 32^. Similarly, tumour cells can rewire their metabolism to suit their needs under different environment conditions, which can negatively affect their response to common antitumoral drugs, such as mTOR^9^ and glutaminase inhibitors^33^, 5-fluorouracil and metformin, and even to immunotherapy^10^. Consequently, the inhibition of this metabolic plasticity is becoming a promising therapeutic target, and its cell selectivity is undoubtedly hugely important in the context of brain tumours. Therefore, here we investigated the effect of TAT-Cx43_266-283_, a promising therapeutic peptide against glioma, on GSC metabolism and metabolic plasticity as a meaningful predictor of the success of a future clinical application.

## Results

### TAT-Cx43_266-283_ decreases glucose uptake selectively in GSCs

Because the highly efficient glucose uptake in GSCs confers these cells with a competitive advantage within the brain environment^13^, we first analysed the effect of TAT-Cx43_266-283_ on glucose transport and uptake in GSCs, neurons and astrocytes. Glucose transport was evaluated in GSCs using the non-metabolizable fluorescent glucose analogue 6-NBDG, whose uptake reflects gradient-driven influx through glucose transporters^34^. We found no differences in 6-NBDG transport (146 μM for 1 h) between controls (untreated condition (Control) and negative control (TAT)) and TAT-Cx43_266-283_-treated GSCs (50 μM, 24 h), suggesting that TAT-Cx43_266-283_ does not modify glucose transport under these experimental conditions (Fig. 1a).

**Figure 1.**
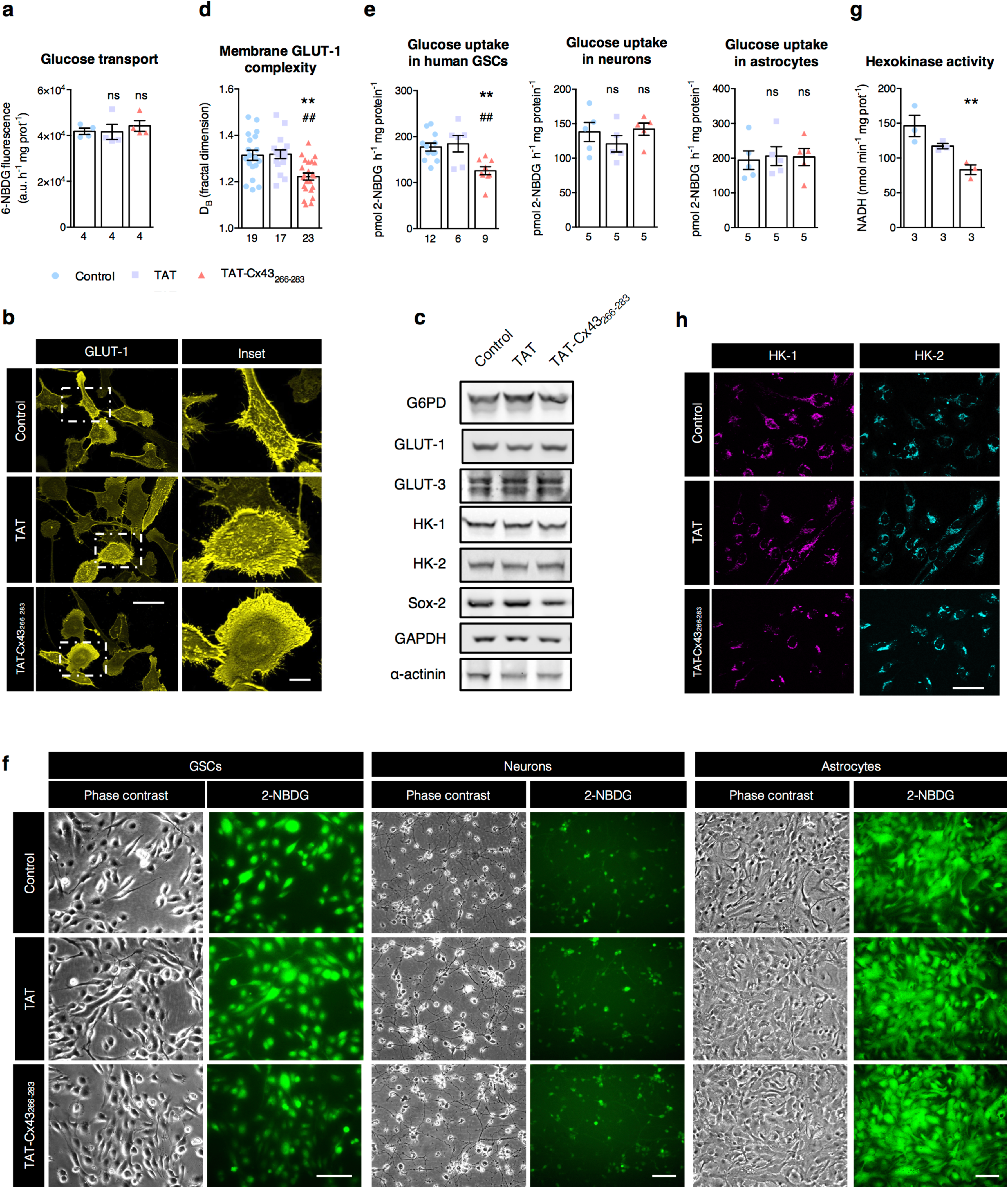
TAT-Cx43_266-283_ decreases glucose uptake through HK activity selectively in GSCs. Cells were treated with 50 μM TAT or TAT-Cx43_266-283_ for 24 h prior to the assays. (**a**) Glucose transport was estimated with the non-phosphorylatable fluorescent glucose analogue 6-NBDG. GSCs were incubated with 146 μM 6-NBDG for 1 h and lysed, and 6-NBDG fluorescence was measured on a microplate fluorometer. Results were normalized to milligrams of protein in each sample. (**b**) Representative micrographs of GSCs immunostained for GLUT-1 showing that TAT-Cx43_266-283_ changes the GLUT-1 staining pattern in the cell membrane without affecting GLUT-1 signal intensity. Scale bar: 50 μm; inset: 10 μm. (**c**) Representative western blots of the indicated proteins. GAPDH and α-actinin blots are shown as loading controls. (**d**) Changes in the GLUT-1 staining pattern analysed by measurement of membrane GLUT-1 complexity as the fractal dimension of the micrographs. (**e**, **f**) Cells were incubated with 146 μM of the fluorescent glucose analogue 2-NBDG for 1 h. (**e**) Representative micrographs showing that, while 2-NBDG uptake was not affected in neurons and astrocytes, GSCs showed reduced 2-NBDG uptake promoted by TAT-Cx43_266-283_. Scale bar: 100 μm. (**f**) Quantification of 2-NBDG uptake in GSCs, neurons and astrocytes. After incubation with 2-NBDG, the cells were lysed and 2-NBDG fluorescence was measured on a microplate fluorometer. The 2-NBDG concentration in the samples was determined using a standard curve. Results were normalized to milligrams of protein in each sample. (**g**) GSCs immunostained for HK-1 (magenta) and HK-2 (cyan). Representative micrographs showing that HK-1 and HK-2 display mostly a mitochondrial distribution and that, in GSCs treated with TAT-Cx43_266-283_, HK-1 and HK-2 staining is condensed near the cell nuclei. Scale bar: 50 μm. (**h**) Hexokinase activity in GSCs determined as described in Methods. The absorbance of the cell lysates was measured on a microplate fluorometer at two time points within the linear range of the reaction and normalized to milligrams of protein in each sample. All data are mean ± s.e.m. and were obtained from at least three independent experiments with at least two technical replicates (one-way ANOVA: \*\**P* < 0.01, \*\*\**P* < 0.001 vs control; ^##^*P* < 0.01, ^###^*P* < 0.001 vs TAT; ns, not significant). Numbers under bars indicate the number of independent experiments (**a**, **e**, **h**) or the number of images analysed (**d**).

To confirm this result, we analysed GLUT-1, the main glucose transporter. GLUT-1 levels in GSCs were not affected by TAT-Cx43_266-283_ treatment (Fig. 1b and c and Supplementary Fig. 1a and b). However, the complexity of the GLUT-1 staining pattern in the cell membrane decreased after TAT-Cx43_266-283_ treatment (Fig. 1b, inset). To quantify this finding, the fractal dimension of the GLUT-1 images was calculated as an index of membrane complexity (see Methods). The lower values obtained for the TAT-Cx43_266-283_ images suggest that TAT-Cx43_266-283_ decreases the complexity of membrane GLUT-1 staining in GSCs (Fig. 1d). This might be due to changes in the polymerization state of the actin cytoskeleton caused by TAT-Cx43_266-283_^35^, which can modulate GLUT-1 clustering without affecting glucose transport^36^. The levels of GLUT-3, another glucose transporter frequently co-opted by cancer cells^13^, were not affected by TAT-Cx43_266-283_ under these experimental conditions (Fig. 1c).

Next, we measured glucose uptake with the non-metabolizable fluorescent glucose analogue 2-NBDG, which is phosphorylated by HK and trapped intracellularly. The results showed that TAT-Cx43_266-283_ decreased glucose uptake in GSCs by ∼30%, whereas uptake was unchanged in neurons and astrocytes (Fig. 1e and f). This result adds to the evidence from our previous report that TAT-Cx43_266-283_ specifically targets GSCs^6^. Moreover, TAT-Cx43_266-283_ reduced HK activity (Fig. 1g) without altering the levels of HK-1 and HK-2 in GSCs treated with TAT-Cx43_266-283_ (Fig. 1c and Supplementary Fig. 1c). Because HK-1 and HK-2 activity is upregulated by c-Src phosphorylation^12^, the TAT-Cx43_266-283_–induced inhibition of c-Src^7, 26^ might hinder HK activity. Immunofluorescence assays revealed that both HKs localized more compactly around the nuclei of GSCs treated with TAT-Cx43_266-283_ (Fig. 1h). Taken together, these data indicate that 24-h treatment with TAT-Cx43_266-283_ reduces glucose uptake selectively in GSCs through decreased HK activity, without affecting glucose transport or the protein levels of GLUT-1, GLUT-3, HK-1 or HK-2.

### TAT-Cx43_266-283_ reduces mitochondrial metabolism without increasing glycolysis in GSCs

Because we found that TAT-Cx43_266-283_ altered the location of HKs (Fig. 1h), which are frequently associated with mitochondria^11, 12^, we used a mitochondrial dye to follow the cellular distribution of these organelles. Importantly, these images revealed that TAT-Cx43_266-283_ treatment (50 μM, 24 h) modified the localization of mitochondria in GSCs but not in astrocytes (Fig. 2a and Supplementary Fig. 2a). Indeed, the distance from the furthest mitochondria to the nucleus was reduced in GSCs after TAT-Cx43_266-283_ treatment (Fig. 2b), suggesting altered mitochondrial trafficking. This change in mitochondrial localization might arise from cytoskeleton rearrangement^35^ and contribute to the reported TAT-Cx43_266-283_–induced inhibition of GSC invasion and migration^26^, as previously shown by others^37^.

**Figure 2.**
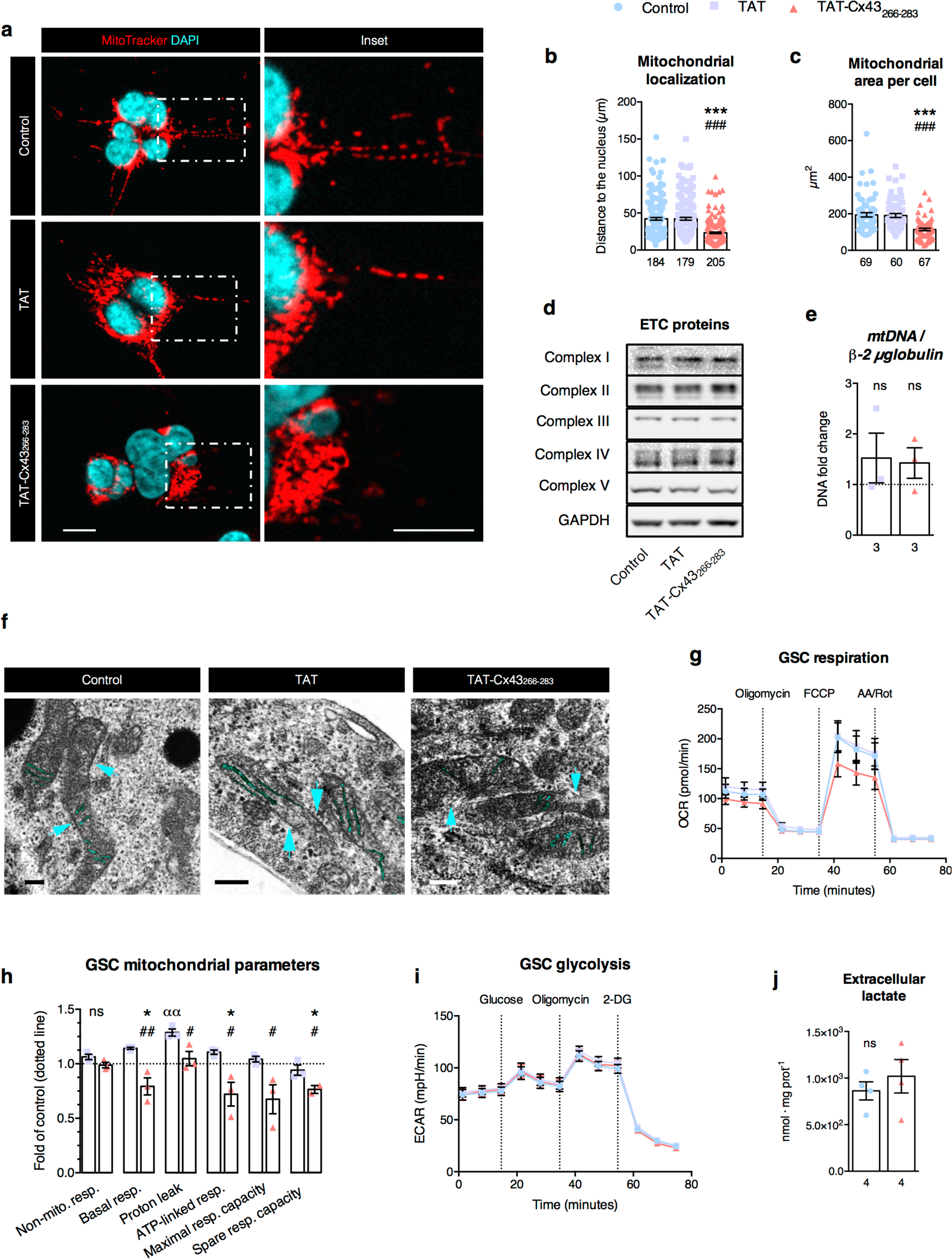
TAT-Cx43_266-283_ induces changes in the localization, ultrastructure and function of mitochondria in GSCs. GSCs were treated with 50 μM TAT or TAT-Cx43_266-283_ for 24 h prior to the assays. (**a**) GSCs were incubated with MitoTracker for 1 h and then fixed, counterstained with DAPI and imaged. Mitochondria in GSCs treated with TAT-Cx43_266-283_ appeared closer to the cell nucleus and more condensed than those of control or TAT-treated GSCs. Scale bar: 10 μm. (**b**) Distance from the furthest mitochondria to the centre of the nucleus in each cell analysed. (**c**) Quantification of mitochondrial area per cell. The mitochondrial network of each cell was selected as a ROI based on MitoTracker staining and the area of each ROI was measured. (**d**) Representative western blots of electron transport chain (ETC) subunit proteins from complexes I (NDUFB8), II (SDHB), III (UQCRC2), IV (MTCO1) and V (ATP5A). No changes were observed in the relative levels of these mitochondrial proteins under the indicated experimental conditions. (**e**) Quantification of mitochondrial DNA relative to nuclear DNA by qPCR. (**f**) GSCs were imaged with transmission electron microscopy. In GSCs treated with TAT-Cx43_266-283_, the mitochondrial outer membrane and cristae were less defined, and the mitochondrial matrix was more electron-dense. Scale bar: 200 nm. (**g**) The OCR of GSCs was measured after addition of 1.5 μM oligomycin to block ATP-linked OCR, 0.4 μM FCCP to uncouple mitochondria to the obtain maximal OCR and 0.5 μM rotenone/antimycin A (Rot/AA) to shut down mitochondrial respiration. Data from three independent experiments. (**h**) Quantification of mitochondrial parameters in GSCs obtained from the data shown in (**g**). (**i**) The ECAR of GSCs was measured after addition of 10 mM glucose to assess the glycolysis rate, 1 μM oligomycin to obtain the maximal ECAR and 50 mM 2-deoxyglucose to shut down glycolysis. Data from four independent experiments. (**j**) Extracellular lactate measured in control and TAT-Cx43_266-283_–treated GSCs. All data are mean ± s.e.m. and were obtained from at least three independent experiments (except for transmission electron microscopy images) with at least two technical replicates (one-way ANOVA: \**P* < 0.05 vs control, \*\**P* < 0.01 vs control; ^#^*P* < 0.05 vs TAT, ^##^*P* < 0.01 vs TAT; ^αα^*P* < 0.01 vs TAT-Cx43, ns, not significant). Numbers under bars indicate the number of cells analysed (**b, c**) or the number of biological replicates (**e, j**).

In addition, the area occupied by mitochondria per cell was reduced in TAT-Cx43_266-283_–treated GSCs (Fig. 2c). To establish if this result is due to the spatial rearrangement of mitochondria or to a decrease in mitochondrial mass, we performed western blotting against subunit proteins from each of the five complexes of the electron transport chain (ETC) involved in oxidative phosphorylation (OXPHOS). Treatment with TAT-Cx43_266-283_ did not affect the relative abundance of ETC complexes in GSCs (Fig. 2d and Supplementary Fig. 2b). Similarly, quantification of mitochondrial DNA relative to nuclear DNA by real-time quantitative polymerase chain reaction (qPCR) did not reveal any changes (Fig. 2e and Supplementary Fig. 2c), confirming that 24-h TAT-Cx43_266-283_ treatment induced a spatial rearrangement in the mitochondrial network of GSCs without changes in total mitochondrial mass. To further characterize the effect of TAT-Cx43_266-283_ on GSC mitochondria, we performed transmission electron microscopy. The images revealed that, in mitochondria from TAT-Cx43_266-283_– treated GSCs, the matrix was more electron-dense and the outer membrane and cristae less defined, indicative of structural changes associated with the previous observations (Fig. 2f; the blue arrowheads indicate the outer membrane, the cristae are coloured).

To determine if the distinct cellular localization and ultrastructure observed in mitochondria from GSCs treated with TAT-Cx43_266-283_ affected mitochondrial function, we measured GSC oxygen consumption rate (OCR) as an indicator of mitochondrial respiration in a Seahorse XF Analyzer^38^. As shown in Fig. 2g, GSCs treated with TAT-Cx43_266-283_ had a lower basal OCR (minutes 0–18), as well as a lower response to mitochondrial stimulation with FCCP (minutes 42– 55). Mitochondrial parameters^38^ calculated from these data revealed that TAT-Cx43_266-283_ decreased basal respiration and ATP-linked respiration, as well as maximal and spare respiratory capacities (Fig. 2h). Phosphorylation of ETC complexes by c-Src regulates respiration and cellular ATP content^39^, which provides a mechanistic explanation for the decreased mitochondrial activity induced by TAT-Cx43_266-283_—a c-Src–inhibiting peptide^7, 26^—, despite unchanged total levels of ETC complexes. Importantly, analysis of GSC glycolysis (Seahorse XF Analyzer^38^) after TAT-Cx43_266-283_ treatment did not reveal any changes in the extracellular acidification rate (ECAR; mostly resulting from lactate production) (Fig. 2i and Supplementary Fig. 2d). We confirmed this result by analysing extracellular lactate in GSCs after TAT-Cx43_266-283_ treatment (Fig. 2j). Recent reports show that drug resistance to standard therapies, such as bevacizumab, relies on the ability of cancer cells to switch their metabolism from OXPHOS to glycolysis^8^. Therefore, it is noteworthy that, despite the well-characterized ability of cancer stem cells to adapt to new metabolic requirements^10, 13, 14^, GSCs treated with TAT-Cx43_266-283_ did not show enhanced glycolysis to compensate for the decreased mitochondrial activity (Fig. 2g vs 2i). In addition, we did not find any significant effect on OCR or ECAR in neurons or astrocytes treated with TAT-Cx43_266-283_ (Supplementary Fig. 2e), in agreement with the cell-specific effect of TAT-Cx43_266-_ ^6^. Taken together, these data indicate that TAT-Cx43_266-283_ specifically downregulates mitochondrial metabolism in GSCs without increasing aerobic glycolysis, which suggests an impairment of metabolic plasticity upon TAT-CX43_266-283_ treatment.

### TAT-Cx43_266-283_ impairs the metabolic plasticity of GSCs

The plasticity of the metabolic phenotype of cancer stem cells is crucial in the ever-changing tumour microenvironment^8, 11, 13, 14^. In order to characterize the ability of the GSCs used in this study to adapt their metabolism to shifts in nutrient availability, we used time-lapse microscopy to analyse their survival in media with different nutrient contents. Interestingly, GSCs survived for 4 days with only glucose or only amino acids in the culture medium (Fig. 3a, Supplementary Fig. 3a and Videos 1–2). Under these starvation conditions, GSCs eventually adopted a dormancy state, yet they were able to resume proliferation when nutrients became available once again (Fig. 3a and videos 1.2 and 2.2), demonstrating a high metabolic plasticity.

**Figure 3.**
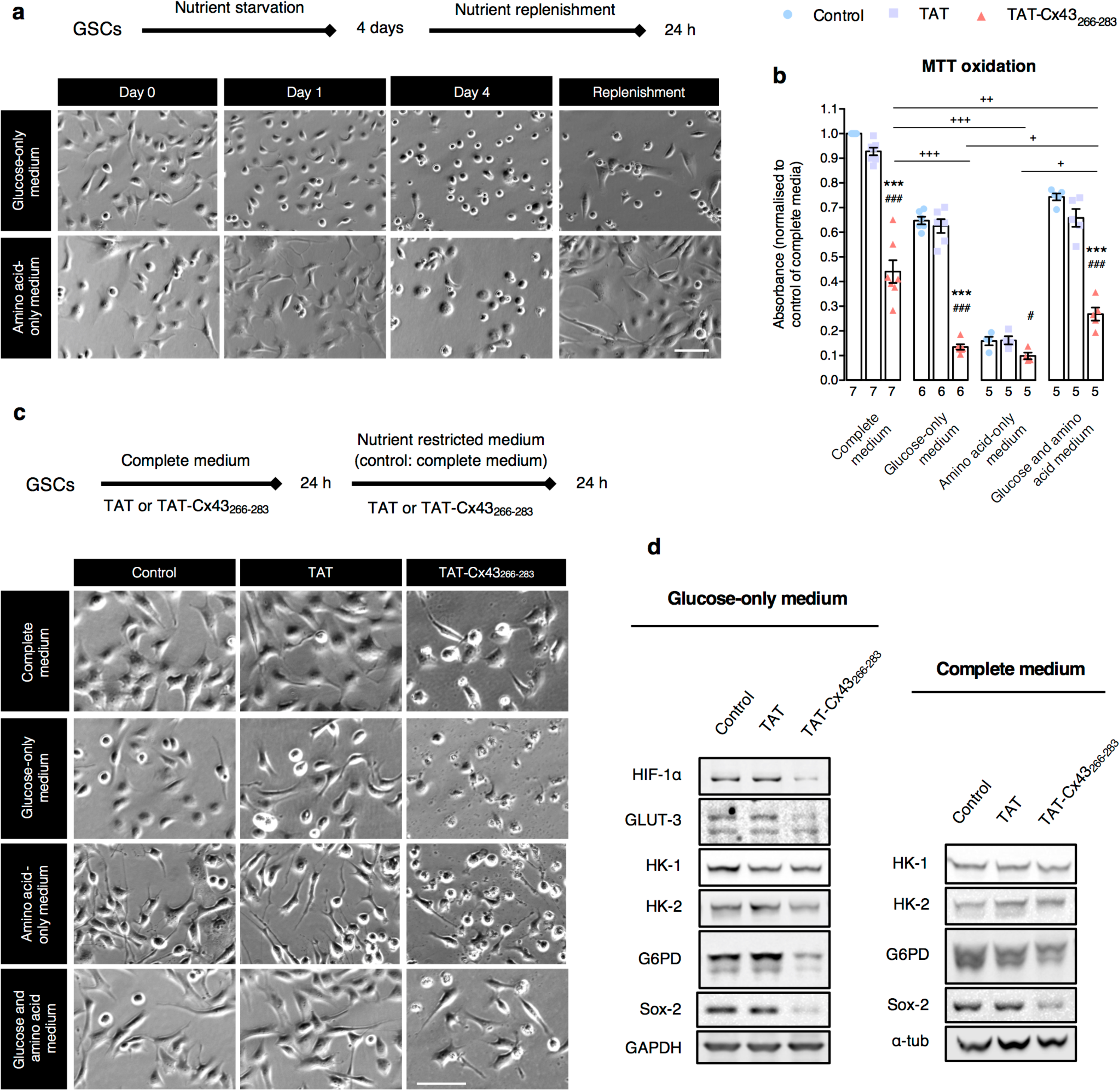
TAT-Cx43_266-283_ maintains its antitumour effect under several nutrient-deprived conditions. (**a**) GSCs were recorded by time-lapse microscopy in medium containing only glucose or only amino acids (Videos 1 and 2, respectively). After 4 days, these media were changed to complete medium and cells were recorded for a further 24 h (nutrient replenishment). Note that cells that appeared dead (bright, round shape) could survive and proliferate after nutrient replenishment, which was not true for cells maintained in medium with no nutrients at all (Supplementary Fig. 3 and Video 3). Scale bar: 100 μm. (**b–e**) GSCs were treated with 50 μM TAT or TAT-Cx43_266-283_ in complete medium for 24 h. Then, the medium was replaced with medium containing glucose (glucose-only medium), amino acids (amino acid-only medium) or both glucose and amino acids, as well as 50 μM of each peptide, for another 24 h. For comparison, cells were treated in parallel with complete medium and 50 μM of peptides. Then, the indicated assays were carried out. (**b**) MTT oxidation of GSCs under the indicated conditions. (**c**) GSCs were recorded by time-lapse microscopy in complete or glucose-only medium (Videos 4–9; the images are the last photomicrograph from each video) or photographed under the indicated conditions. Scale bar: 100 μm. (**d**) Representative western blots of the indicated proteins under the indicated conditions. GSCs treated with TAT-Cx43_266-283_ in glucose-only medium exhibit decreased levels of HIF-1α, GLUT-3, HK-2, G6PD and Sox-2, but not HK-1 (right panel). However, GSCs treated in complete medium showed only slight decreases (left panel). GAPDH and α-tubulin blots are shown as loading controls. All data are mean ± s.e.m. and were obtained from at least three independent experiments with at least two technical replicates (one-way ANOVA: \*\*\**P* < 0.001 vs control; _###_*P* < 0.001 vs TAT; ^+^*P* < 0.05, ^++^*P* < 0.01, ^+++^*P* < 0.001 between the indicated conditions). Numbers under bars indicate the number of biological replicates in (**b**).

Emerging data suggest that, beyond cell-intrinsic factors, nutrient availability in the tumour microenvironment can also influence drug response, hampering the efficacy of common antitumoral drugs^10^. To investigate whether this was also the case for TAT-Cx43_266-283_, GSCs in complete medium were treated with 50 μM TAT or TAT-Cx43_266-283_ for 24 h and then switched to glucose-only, amino acid-only or glucose and amino acid media containing 50 μM TAT or TAT-Cx43_266-283_ for a further 24 h. For comparison, GSCs were cultured in parallel in complete medium. Time-lapse microscopy and MTT assays revealed that TAT-Cx43_266-283_ maintained its antitumour effect under the different nutrient-deprived conditions (Fig. 3b and c and Videos 4–9). Consistent with the TAT-Cx43_266-283_–induced reduction in glucose uptake and metabolism, we found a pronounced increase in cell death in GSCs treated with TAT-Cx43_266-283_ in glucose-only medium compared with those cultured in any other medium (Fig. 3b and c and Videos 4– 9).

At the molecular level, GSCs adapted to glucose-only medium by upregulating glucose metabolism enzymes critical for cancer cells, such as HK-2^11, 12^ and glucose-6-phosphate-dehydrogenase (G6PD)^40, 41^, the key and rate-limiting enzyme in the pentose phosphate pathway (Supplementary Fig. 3b). However, GSCs treated with TAT-Cx43_266-283_ in glucose-only medium showed markedly decreased levels of both HK-2 and G6PD, as well as GLUT-3, and, importantly, HIF-1α, a master regulator of glucose metabolism^42, 43^(Fig. 3d). Meanwhile, HK-1 levels remained unchanged, suggesting that this is not a general reduction in glucose metabolism (Fig. 3d). In addition, our results confirm that TAT-Cx43_266-283_ targets the metabolic adaptation to this nutrient-deprived condition because GSCs in complete medium showed no or only slight decreases in the levels of these proteins upon treatment with TAT-Cx43_266-283_ for 24 h or 48 h, respectively (Fig. 1e and Fig. 3d). On a different note, Sox-2, a transcription factor implicated in stem cell maintenance^7^, was also up-regulated in GSCs cultured in glucose-only medium (Supplementary Fig. 3b). This is in agreement with previous studies showing cancer stem cell enrichment under nutrient-deprived conditions and the role of certain master stem cell transcription factors in promoting preferential glucose metabolism^13, 44^. According to our previous results^7^, Sox-2 levels in treated GSCs were decreased in complete medium (Fig. 3d). Interestingly, here we found that this reduction also takes place in glucose-only medium, suggesting TAT-Cx43_266-283_ abrogates GSC enrichment under this condition (Fig. 3d). Together, these results indicate that GSCs treated with TAT-Cx43_266-283_ fail to upregulate proteins necessary for survival in different metabolic scenarios, particularly those needed to exploit glucose as an energy source in the absence of other nutrients (e.g., amino acids/proteins or lipids) (Fig. 3d and Supplementary Fig. 3b). This lack of adaptation might be responsible for maintaining the effect of TAT-Cx43_266-283_ in different metabolic environments, in contrast to other antitumour therapies^8, 10^.

### TAT-Cx43_266-283_ impairs glucose metabolism ex vivo and in vivo

To explore the effect of TAT-Cx43_266-283_ on glucose metabolism in GSCs within the brain environment, we first evaluated glucose uptake in an ex vivo model of glioma^6^. As illustrated in Fig. 4a, fluorescently labelled GSCs were placed in organotypic brain slices and allowed to engraft overnight. The GSC–organotypic brain slice co-cultures were treated with 50 μM TAT or TAT-Cx43_266-283_ for 48 h, incubated with 146 μM 2-NBDG in glucose-free medium for 1 h and mounted for microscopy (Fig. 4a and Supplementary Fig. 4a). In agreement with the in vitro 2-NBDG study (Fig. 1e and f), glucose uptake was reduced to ∼65% in GSCs in co-cultures treated with TAT-Cx43_266-283_ compared with the control and TAT conditions (Fig. 4b and c), with no apparent effect on the brain parenchyma (Fig. 4c and Supplementary Fig. 4a).

**Figure 4.**
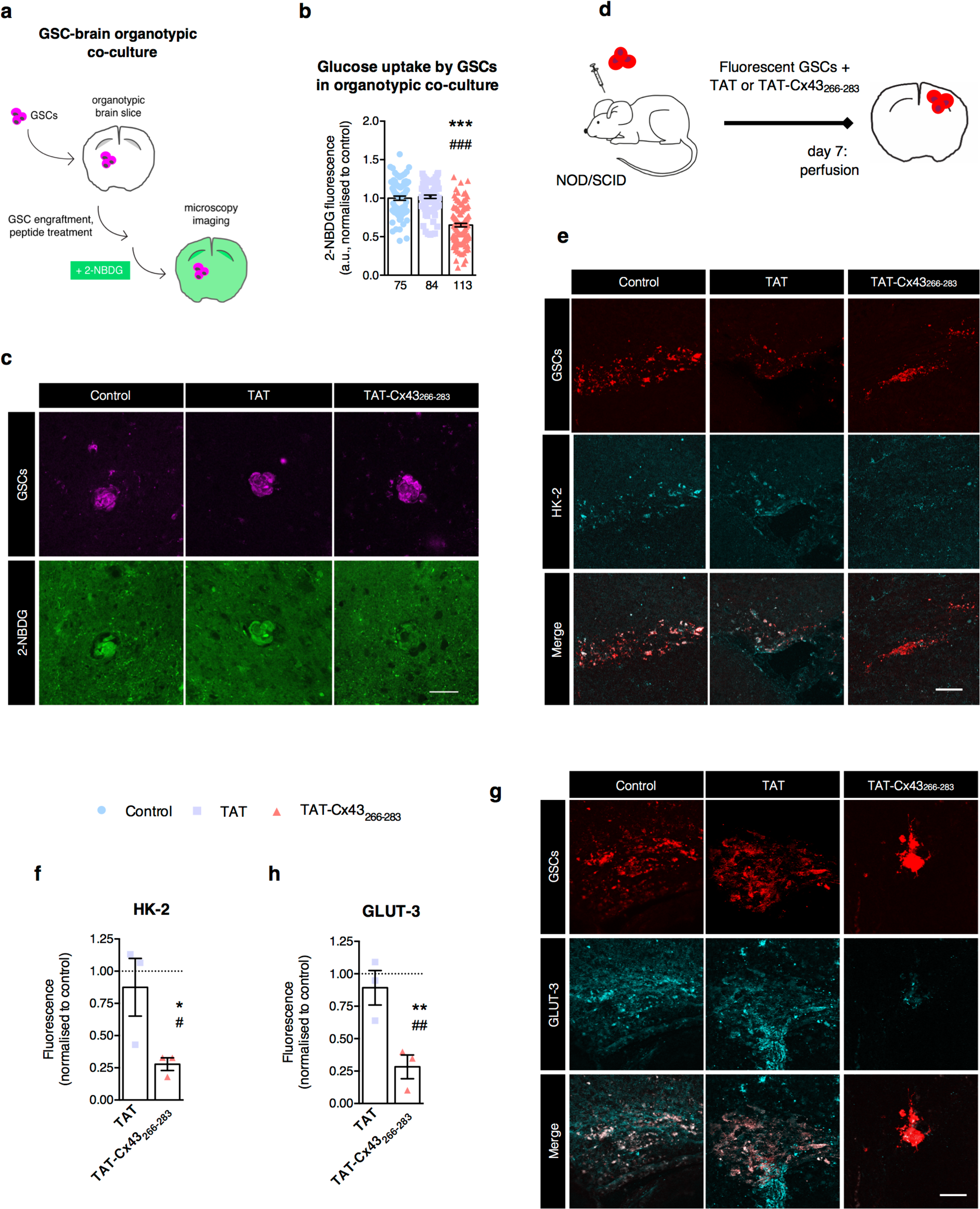
TAT-Cx43_266-283_ impairs glucose metabolism ex vivo and in vivo. (**a**) Schematic drawing of the 2-NBDG uptake experiments in GSC–organotypic brain slice co-cultures. 2,500 fluorescently labelled GSCs were placed onto each organotypic brain slice and allowed to integrate overnight. The GSC–brain slice co-cultures were incubated with 50 μM TAT or TAT-CX43_266-283_ for 48 h. Then, the co-cultures were incubated with 146 μM 2-NBDG for 1 h and analysed by confocal microscopy. (**b**) 2-NBDG fluorescence (mean grey value) was quantified in the regions of interest (ROIs) corresponding to GSC fluorescence in confocal microscopy images. (**c**) Representative confocal microscopy images showing the reduction in 2-NBDG uptake (green) specifically in GSCs (magenta) in the GSC–brain slice co-cultures treated with TAT-Cx43_266-283_. Scale bar: 50 μm. (**d**) Schematic drawing of the in vivo experimental design. Briefly, 5,000 human GSCs fluorescently labelled with PKH26 or expressing mCherry were intracranially injected alone (control) or together with 100 μM TAT or TAT-Cx43_266-283_ into the brain of NOD/SCID mice. After 7 days, the animals were perfused and their brains processed for immunofluorescence and analysed by confocal microscopy. (**e** and **g**) Representative images of control, TAT and TAT-Cx43_266-283_ brain sections showing GSCs (red), HK-2 (**e**) and GLUT-3 (**f**) and merged images of the same fields. For DAPI staining and the specificity of antibodies against human HK-2 and GLUT-3, see Supplementary Fig. 4. Scale bar: 50 μm. (**f** and **h**) Quantification of HK-2 (**f**) and GLUT-3 (**h**) fluorescence in GSCs in brain sections of a xenograft mouse model of glioma. Mean fluorescence intensity was normalized to control (assigned a value of 1; dotted line). Three animals were analysed per condition. Data are mean ± s.e.m. and were obtained from at least three independent experiments (one-way ANOVA: \**P* < 0.05 vs control, \*\**P* < 0.01 vs control, \*\*\**P* < 0.001 vs control; ^#^*P* < 0.05 vs TAT, ^##^*P* < 0.01 vs TAT, ^###^*P* < 0.001 vs TAT). Numbers under bars indicate the number of ROIs analysed (**b**).

Next, we used an in vivo glioma model consisting of the intracranial injection of 5,000 fluorescently labelled human GSCs together with 100 μM TAT or TAT-Cx43_266-283_ into the brains of NOD/SCID mice^6^. Seven days after surgery, the mice were sacrificed, and their brains were processed for analysis (Fig. 4d). We had already found that TAT-Cx43_266-283_ in vitro impaired HK activity after 24 h of treatment (Fig. 1g) and decreased HK-2 levels after 48 h of treatment in glucose-only medium (Fig. 3d). We corroborated these findings in vivo through immunofluorescence analysis of HK-2 in the GSCs of the xenografted mice, who showed a ∼75% reduction in HK-2 staining in GSCs injected with TAT-Cx43_266-283_ (Fig. 4e and f and Supplementary Fig. 4b and d).

Within the bulk glioma, GLUT-3 is preferentially expressed by GSCs in vivo and its inhibition impairs GSC growth and tumorigenicity, emerging as an important target in malignant gliomas^13^. Because our in vitro results showed a decreased level of GLUT-3 in glucose-only medium (Fig. 3d), we analysed its level in the in vivo implanted GSCs as well. Notably, immunofluorescence analysis revealed a ∼75% reduction in GLUT-3 staining in GSCs injected with TAT-Cx43_266-283_ (Fig. 4g and h and Supplementary Fig. 4c and e). This finding and the reduced HK-2 levels in GSCs in vivo recapitulate the HK-2 and GLUT-3 decrease found in GSC treated in glucose-only medium in vitro (Fig. 3d).

## Discussion

Our previous studies showed that GSCs can be specifically targeted by a short cell-penetrating peptide based on Cx43 (TAT-Cx43_266-283_) that reduces tumour growth in preclinical models^6^ via c-Src inhibition^7^. The effects of TAT-Cx43_266-283_ on GSC metabolism found in this study highlight the therapeutic potential of this compound against glioblastoma because TAT-Cx43_266-283_ targets crucial proteins for glioma malignancy, namely GLUT-3, HK-2, G6PD and HIF-1α^11–13, 40–42^. Our in vitro, ex vivo and in vivo data show that glucose uptake is specifically impaired in GSCs through short- and long-term mechanisms involving the regulation of the activity and expression of key glucose enzymes. Indeed, our results suggest an initial short-term regulation of HK activity, presumably through the inhibition of its phosphorylation by c-Src^12^. This is followed by long-term regulation through reduced HK-2 protein levels, which is highly relevant considering the strong link between HK activity, particularly that of HK-2, and the malignant phenotype^11, 12^. We hypothesize that, because c-Src activity stabilizes the levels of HIF-1α ^29, 30^, TAT-Cx43_266-283_ might decrease HIF-1α through c-Src inhibition, leading to the downstream downregulation of HK-2, G6PD and GLUT-3 shown here. In agreement with these results, Cx43 has been proposed to regulate glucose uptake, HK-2 and GLUT-3 through c-Src and HIF-1α^29^. In addition, TAT-Cx43_266-283_ increases the levels of PTEN^26^, which could also contribute to blocking HIF-1α-dependent gene transcription^45^.

GLUT-3 has been pinpointed as an excellent target to reduce glioma progression because of the high GSC dependence on this glucose transporter^13^. However, in the brain context, the inhibition of GLUT-3 is challenging because it is also the main glucose transporter in neurons^46^. Remarkably, we found that TAT-Cx43_266-283_ downregulates GLUT-3 in GSCs without affecting glucose uptake in neurons, probably because its regulation in neurons is independent of c-Src activity^46^. In fact, TAT-Cx43_266-283_ appears to be innocuous in terms of neuron and astrocyte metabolism, presumably because of the low basal activity of c-Src in healthy cells^7^.

It has been reported that GSCs use OXPHOS as their primary energy source, for which they require an intact mitochondrial respiratory chain to survive and sustain their tumorigenic potential^47^. As previously mentioned, c-Src, via phosphorylation of ETC complexes, regulates mitochondrial respiration necessary for the survival of glioma cells^39^. Consistent with these reports, here we show that the c-Src inhibiting peptide, TAT-Cx43_266-283,_ impairs glucose mitochondrial metabolism and promotes GSC death. Moreover, ATP generated in the ETC is necessary for the c-Src-activating phosphorylation at Y416^48^, which might create a negative regulatory loop when ATP becomes scarce due to TAT-Cx43_266-283_ treatment. In terms of glioma therapy, the impairment of mitochondrial metabolism is highly relevant because it might per se increase the effectiveness of radiotherapy^4^. Importantly, the reduced mitochondrial activity promoted by TAT-Cx43_266-283_ is not compensated by an increase in glycolysis, which is critical to avoid drug resistance^8–10^. Indeed, TAT-Cx43_266-283_ exerts antitumour effects in different nutrient-depleted media, where it prevents metabolic plasticity and eventually leads to cell death. This is especially relevant considering the growing body of work acknowledging intrinsic and extrinsic metabolic states as key factors in cancer drug resistance^8, 10, 32, 49^.

Although further research is needed to address the role of TAT-Cx43_266-283_ in amino acid, lipid and protein metabolism, our in vivo work shows that the effects of TAT-Cx43_266-283_ on GSC metabolism extend beyond in vitro conditions, indicating that they are independent of the metabolic tumour microenvironment, contrary to other antitumoral drugs^10^. The metabolically challenging tumour environment promotes GSC enrichment^13, 15, 16^, however, TAT-Cx43_266-283_ reduces GLUT-3 and HK-2 levels in GSCs in vivo, two proteins involved in glioma establishment, maintenance and resistance^11–13^. Concomitantly, in the same glioma model used in this study, TAT-Cx43_266-283_ induces the loss of the stemness phenotype of the implanted GSCs, as evidenced by decreased expression of Sox-2 and nestin in these cells 7 days post-implantation, and impairs glioma growth 30 days post-implantation, leading to decreased tumorigenicity and increased survival^6^. Given our in vitro, ex vivo and in vivo results and the fundamental role played by HK-2^11, 12^ and GLUT-3^13^ in glioma formation and development, TAT-Cx43_266-283_ might impair in vivo glioma growth through both the loss of the stemness phenotype^6^ and the abrogation of metabolic plasticity, two phenomena that are highly dynamic and tightly linked^5, 13–16^.

From a clinal point of view, the search for therapeutic molecules that specifically target the metabolism of cancer stem cells is on the rise^50^, with metabolic plasticity and nutrient composition of the tumour microenvironment at the centre of drug response^10^. Taken together, these results support the notion that metabolic plasticity, rather than specific metabolic pathways, constitutes a major targetable vulnerability^14^, and confirm the value of TAT-Cx43_266-283_ for glioma therapy alone or in combination with therapies whose resistance relies on metabolic adaptation.

## Methods

### Animals

Albino Wistar rats and NOD SCID mice were obtained from the animal facility of the University of Salamanca. All animal procedures were approved by the ethics committee of the University of Salamanca and the *Junta de Castilla y León* and were carried out in accordance with European Community Council Directives (2010/63/UE) and Spanish law (R.D. 53/2013 BOE 34/11370–420, 2013) for the care and use of laboratory animals.

Mice were kept in 12 h light/dark cycles, had free access to food and water at all times and were monitored for signals of humane endpoint daily, including changes in behaviour and weight. A total of nine male and female adult mouse were used for the in vivo experiments.

### Cell cultures

G166 human GSCs (IDH-wt)^51^ were obtained from BioRep and cultured as previously described^7^. The cells were cultured in RHB-A medium (Takara Bio Inc., Condalab) supplemented with 2% B27 (Life Technologies), 1% N2 (Life Technologies), 20 ng ml^-1^ EGF and 20 ng ml^-1^ b-FGF (PeproTech) under adherent conditions, as described by Pollard et al.^52^ (complete medium). Culture plates were coated with 10 μg ml^-1^ laminin (Invitrogen, 23017-015) for 2 h before use. Cells were grown to confluence, dissociated using Accutase (Thermo Fisher) and then split to convenience. We routinely used cultures expanded for no more than 15 passages.

Primary rat neuron and astrocyte cultures were carried out as previously described^53^. Neurons in primary culture were prepared from the forebrains of foetuses at 17.5 days of gestation and cultured in Dulbecco’s modified Eagle’s medium (DMEM; Sigma-Aldrich, D5523) supplemented with 10% foetal calf serum (FCS; Gibco). One day after plating, cytosine arabinoside was added to avoid glial cell proliferation. Astrocytes in primary culture were prepared from the forebrains of 1- to 2-day-old Wistar rats and cultured in DMEM supplemented with 10% FCS. The cells were maintained at 37 °C in an atmosphere of 95% air/5% CO_2_ and with 90–95% humidity.

### GSC–organotypic brain slice co-cultures

Organotypic brain slice cultures were prepared as previously described^54^. Briefly, 350-μm-thick brain slices were obtained from neonatal Wistar rats and cultured onto cell culture inserts in DMEM supplemented with 10% horse serum and glucose (final concentration, 33 mM). The medium was replaced three times a week and slices were maintained in culture for 19–20 DIV. Then, 2,500 G166 GSCs fluorescently labelled with CellTracker Red CMPTX (Life Technologies) were placed onto each brain slice and maintained in culture for the indicated time.

### Cell treatments

Synthetic peptides (> 85% pure) were obtained from GenScript (Piscataway, NJ, USA). YGRKKRRQRRR was used as the TAT sequence, which enables the cell penetration of the peptides^55^. The TAT-Cx43_266-283_ sequence was TAT-AYFNGCSSPTAPLSPMSP (patent ID: WO2014191608A1).

For in vitro and ex vivo studies, the peptides (TAT, as a negative control, and TAT-Cx43_266-283_) were used at 50 μM in culture medium at 37 °C for 24 and 48 h, respectively. For in vivo studies in NOD SCID mice, a single intracranial injection of 1 μl of saline containing 100 μM TAT or 100 μM TAT-Cx43_266-283_ and 5,000 GSCs was performed.

In addition to either 50 μM TAT or TAT-Cx43_266-283_ (where indicated), the cell media (pH 7.3) used in the metabolic plasticity study had the following compositions: in Video 1, Earl’s Balanced Salt Solution (EBSS) containing 5.6 mM glucose; in Video 2, EBSS containing 0.4 mM glycine, 4 mM glutamine, 4 mM serine and essential amino acids (MEM Amino Acids Solution; Gibco, 11130036); for all other experiments, Earl’s Balanced Salt Solution (EBSS) containing 14 mM glucose (glucose-only medium); EBSS containing 4 mM glutamine, essential amino acids and non-essential amino acids (MEM Non-Essential Amino Acids Solution; Gibco, 11140050) (amino acid-only medium); and EBSS containing 14 mM glucose, 4 mM glutamine, essential amino acids and non-essential amino acids (glucose and amino acid medium).

### In vitro 2-NBDG and 6-NBDG uptake assay

Neurons were seeded at 10^5^ cells/cm^2^ in 35 mm culture dishes and grown for 3 days in vitro before incubation with the peptides. Astrocytes were seeded at 1.3 × 10^4^ cells/cm^2^ and incubated with the peptides when confluence was reached after 19–21 DIV. GSCs cells were seeded at 6 × 10^4^ cells/cm^2^ and allowed to attach overnight before incubation with the peptides. Astrocytes and GSCs cells were seeded in 12-well plates. The cells were incubated for 30 min in glucose-free medium (RPMI, Sigma-Aldrich) and subsequently for 1 h in glucose-free medium supplemented with 146 μM of the fluorescent glucose analogue 2-NBDG (2-[N-(7-nitrobenz-2-oxa-1,3-diazol-4-yl) amino]-2-deoxy-D-glucose), which is composed of a glucose moiety with an N-nitrobenzoxadiazole (NBD)-amino group (fluorophore) at carbon 2 replacing the hydroxyl group, or 6-NBDG (6-[N-(7-nitrobenz-2-oxa-1,3-diazol-4-yl) amino]-6-deoxy-D-glucose), whose NBD-amino group is placed at carbon 6 and can therefore not be phosphorylated by HKs^34^ (Thermo Fisher). The cells were then washed with ice-cold phosphate-buffered saline (PBS) and lysed, scraped and homogenized by 10 passages through a 25-gauge needle. Homogenates were centrifuged and the fluorescence of supernatants was measured on a microplate reader (Appliskan; Thermo Electron Corporation, Thermo Scientific). For 2-NBDG uptake, a standard curve was generated by measuring the fluorescence of a range of 2-NBDG concentrations in lysis buffer (1% Nonidet P-40, 1% sodium deoxycholate, 40 mM KCl and 20 mM Trizma Base [Sigma-Aldrich], pH 7.4). 2-NBDG and 6-NBDG uptake was normalized to milligrams of protein measured in the samples.

### Immunofluorescence

For in vitro studies, immunofluorescence was performed as previously described^7^. Briefly, cells were incubated with MitoTracker Red CMXRos (final concentration, 100 nM in culture medium; Invitrogen) for 1 h at 37 °C where indicated and then fixed in methanol for 10 min at –20 °C. The cells were then rinsed in PBS and incubated for 1 h in blocking solution (PBS containing 10% FCS, 0.1 M lysine and 0.02% azide). The samples were incubated overnight at 4 °C with the indicated primary antibody prepared in blocking/permeabilization solution (with 0.1% Triton X-100): rabbit polyclonal antibody against GLUT-1 (1:250; Millipore, 07-1401), rabbit monoclonal antibody against HK-1 (1:200; Cell Signaling Technology, 2024) or mouse monoclonal antibody against HK-2 (1:100; Thermo Fisher, MA5-15679). After repeated washes, they were incubated for 75 min with the corresponding secondary antibody prepared in blocking/permeabilization solution: anti-rabbit IgG or anti-mouse IgG Alexa Fluor 488-, Alexa Fluor 594- or Alexa Fluor 647-conjugated antibodies (1:1,000; Life Technologies). Finally, nuclear DNA was stained with 1 μg ml^-1^ 4’,6-diamidino-2-phenylindole (DAPI) for 1 min. Cells were mounted using a SlowFade Light antifade kit (Life Technologies) and imaged on an inverted Zeiss Axio Observer Z1 microscope for Live-Cell Imaging (Carl Zeiss Microscopy) coupled to an AxioCam MRm camera and Zeiss Apotome (optical sectioning structured illumination microscopy; https://www.zeiss.com/microscopy/int/solutions/reference/all-tutorials/optical-sectioning/apotome-operation.html).

For in vivo studies, sections were washed in PBS and incubated for 1 h in blocking/permeabilization solution. Then, the sections were incubated overnight at room temperature with the indicated primary antibody prepared in blocking/permeabilization solution: mouse monoclonal antibody against HK-2 (1:100; Thermo Fisher, MA5-15679) or mouse monoclonal antibody against GLUT-3 (1:100; Santa Cruz Biotechnology, sc-74497). After repeated washes, the sections were incubated for 2 h with the corresponding secondary antibody prepared in blocking/permeabilization solution: anti-rabbit IgG or anti-mouse IgG Alexa Fluor 488- or Alexa Fluor 647-conjugated antibodies (1:500; Life Technologies). Finally, nuclear DNA was stained with DAPI for 4 min and samples were imaged by confocal microscopy in a Leica TCS SP2 microscope. Briefly, 8-bit images from consecutive focal planes (1 μm along the z axis) were scanned with a pinhole aperture of 1 Airy Unit.

### Image analysis

Images were analysed using Fiji Software^56^, available at http://rsbweb.nih.gov/ij/. For in vitro and ex vivo experiments, details of the image analysis are specified in the corresponding figure legend. In the in vivo study, maximum z projections were obtained from confocal stack images. Projections were subjected to background subtraction and the mean grey value of the image was obtained. Non-specific staining was measured in three identically sized regions from each image and subtracted from the corresponding mean grey value. The fractal dimension of the images was analysed with the FracLac plugin in Fiji (https://imagej.nih.gov/ij/plugins/fraclac/FLHelp/Introduction.htm).

### Hexokinase activity assay

The HK activity of cells was determined using a commercial kit (Abcam, ab136957) following the manufacturer’s instructions. Results were normalized to the protein content of the samples.

### Real-time quantitative polymerase chain reaction

DNA from GSCs was extracted with a QIAamp DNA Mini kit (Qiagen) following the manufacturer’s instructions. DNA concentration and quality were analysed in a NanoDrop 2000 spectrophotometer.

Real-time PCR reactions were performed in triplicate in 96-well plates in a QuantStudio 7 Flex Real-Time PCR System (Thermo Fisher). Equal initial amounts of total DNA were amplified in all conditions. A PCR master mix was prepared for each sample containing 1 μL DNA, 1 μM of forward and reverse primers and 12.5 μL of SYBR® Green Master Mix (Life Technologies) in a 25-μL reaction mix.

The primers used for amplification were: *mt-ND1* (mitochondrial DNA) forward 5’-CCC TAA AAC CCG CCA CAT CT-3’ and reverse 5’-GAG CGA TGG TGA GAG CTA AGG T-3’; *β-2-μglobulin*^57^ (nuclear DNA gene) forward 5’-TGC TGT CTC CAT GTT TGA TGT ATC T-3’ and reverse 5’-TCT CTG CTC CCC ACC TCT AAG T-3’. Negative control reactions for each set of primers were performed in the absence of cDNA template. Reaction products were run on a 2% agarose gel containing Syber Safe DNA gel stain (Invitrogen) to ascertain correct primer amplification (Supplementary Fig. 2).

Real-time PCR results were analysed as described by Pfaffl et al.^58^

### Western blotting

Western blotting was performed as described previously^26^. Briefly, equal amounts of proteins across conditions were separated on NuPAGE Novex Bis-Tris 4–12% Midi gels (Life Technologies) at room temperature and constant voltage. Proteins were transferred to a nitrocellulose membrane (iBlot Gel Transfer Stacks Nitrocellulose) using an iBlot dry blotting system (Life Technologies). After blocking, the membranes were incubated overnight at 4 °C with primary antibodies: mouse monoclonal antibody against α-actinin (1:1,000; Millipore, MAB1682), mouse monoclonal antibody against G6PD (1:250; Santa Cruz Biotechnology, sc-373886), mouse monoclonal antibody against glyceraldehyde phosphate dehydrogenase (GAPDH; 1:5.000; Invitrogen, AM4300), rabbit polyclonal antibody against GLUT-1 (1:500–1:1.000; Millipore, 07-1401), mouse monoclonal antibody against GLUT-3 (1:100; Santa Cruz Biotechnology, sc-74497), rabbit polyclonal antibody against HIF-1α (1:200; Novus Biologicals, NB100-479), rabbit monoclonal antibody against HK-1 (1:250; Cell Signaling Technology, 2024), mouse monoclonal antibody against HK-2 (1:500; Thermo Fisher, MA5-15679), mouse monoclonal antibody against RPL-19 (1:200; Santa Cruz Biotechnology, sc-100830), rabbit polyclonal antibody against Sox-2 (1:500; Abcam, ab97959), Total OXPHOS Rodent WB Antibody Cocktail against NDUFB8 (complex I), SDHB (complex II), UQCRC2 (complex III), MTCO1 (complex IV) and ATP5A (complex V) (1:250; Abcam, ab110413), and mouse monoclonal antibody against α-tubulin (1:1,000; Sigma-Aldrich, T9026). After extensive washing, the membranes were incubated with peroxidase-conjugated anti-rabbit IgG or anti-mouse IgG antibodies (1:5,000; Jackson ImmunoResearch) and developed with a chemiluminescent substrate (Western Blotting Luminol Reagent; Santa Cruz Biotechnology) in a MicroChemi imaging system (Bioimaging Systems). Original and replicate blots are shown in the Supplementary Figures.

### Transmission electron microscopy

Cell culture preparations were fixed in 2% formaldehyde and 2% glutaraldehyde in phosphate buffer for 30 min at 4 °C. Samples were then post-fixed with 1% osmium tetroxide in water, dehydrated through a graded ethanol series and embedded in Epoxy EMbed-812 resin (Electron Microscopy Sciences). Ultrathin sections were obtained with a Leica EM UC7 ultramicrotome, contrasted with uranyl acetate and lead citrate and analysed using a Tecnai Spirit Twin 120 kV electron microscope with a CCD Gatan Orius SC200D camera with DigitalMicrograph™ software. Procedures were performed at the Electron Microscopy Facilities-NUCLEUS of the University of Salamanca.

### Oxygen consumption rate and extracellular acidification rate

The OCR and ECAR were monitored as indicators of mitochondrial respiration and glycolytic function with an XF96 Extracellular Flux Analyzer using an XF Cell Mito Stress Test kit and XF Glycolysis Stress kit according to the manufacturer’s instructions (Seahorse Biosciences). Seeding numbers were optimized to 1.6 × 10^4^ cells/well and 8 × 10^3^ cells/well for GSCs, to 6.36 × 10^4^ and 3.18 × 10^4^ cells/well for neurons and to 2.65 × 10^4^ and 9.93 × 10^3^ cells/well for astrocytes for the Mito Stress Test and Glyco Stress Test kits, respectively.

For the mitochondrial stress test, cells were plated in XF96 plates and treated with TAT or TAT-Cx43_266-283_ as indicated. Prior to the assay, the regular culture media was replaced with Base media (Seahorse Bioscience) supplemented with 1 mM pyruvate, 2 mM glutamine and 10 mM glucose and cells were incubated without CO_2_ for 1 h. Basal rate measurements were taken and then mitochondrial respiratory chain drugs were added, according to Mito Stress kit specifications. 1.5 μM oligomycin was used to block ATP-linked oxygen consumption, 0.4 μM carbonyl cyanide-*P*-trifluoromethoxyphenylhydrazone (FCCP; an uncoupling agent) was used to obtain maximal respiration and 0.5 μM rotenone/antimycin A was used to inhibit complexes I and III to stop all mitochondrial respiration.

For glycolysis analysis, Base media supplemented with 0.5 mM pyruvate and 2 mM glutamine was used and cells were incubated without CO_2_ for 1 h. In accordance with Glycolysis Stress kit specifications, 10 mM glucose was injected to stimulate glycolysis, 1.5 μM oligomycin was then injected to obtain the maximal glycolytic capacity upon oxygen consumption inhibition and finally 50 mM 2-deoxy-D-glucose (2-DG) was used to shut down all glycolysis.

OCR and ECAR were measured three times after the injection of each drug. At least six replicates per condition were performed in each experiment. Parameter calculations were performed using the Seahorse XF Cell Mito Stress Test Report Generator provided by Seahorse Biosciences. Non-mitochondrial respiration is the minimum rate measurement after Rot/AA injection; basal respiration is oxygen consumption used to meet cellular ATP demand, calculated by subtracting non-mitochondrial OCR from the measurement prior to oligomycin addition; proton leak is calculated by subtracting non-mitochondrial OCR from the minimum rate measurement after oligomycin injection; ATP-linked respiration is calculated by subtracting the minimum rate measurement after oligomycin injection from the last rate measurement before oligomycin injection; maximal respiration capacity is calculated by subtracting non-mitochondrial respiration from the maximum rate measurement after FCCP injection; and spare respiratory capacity, the capability to respond to an energetic demand, is calculated as the difference between maximal and basal respiration. Non-glycolytic acidification is the last rate measurement prior to glucose injection, glycolysis is calculated as the maximum rate measurement before oligomycin injection minus the last rate measurement before glucose injection, glycolytic capacity is calculated as the maximum rate measurement after oligomycin injection minus the last rate measurement before glucose injection and glycolytic reserve is calculated as glycolytic capacity minus glycolysis.

### Extracellular lactate

Extracellular lactate was determined using a commercial kit (Sigma-Aldrich, mak064) and the cell media was filtered through 10,000 Nominal Molecular Weight Limit (NMWL) filters to eliminate possible extracellular lactate dehydrogenase that would degrade lactate, following the manufacturer’s instructions. Results were normalized to the protein content of the samples.

### Time-lapse microscopy

GSCs were plated at 5 × 10^4^ cells/well in 12-well plates. Once the cells were attached, TAT or TAT-Cx43_266-283_ was added at 50 μM for 24 h. Then, the cell culture media was changed to the indicated media including 50 μM TAT or TAT-Cx43_266-283_ and the cells were allowed to equilibrate for 1 h in the microscope incubator before imaging. The cells were recorded by time-lapse live-cell imaging for 24 h. Every 10 min, phase-contrast microphotographs of each experimental condition were taken for live-cell imaging with an inverted Zeiss Axio Observer Z1 microscope coupled to an AxioCam MRm camera. The system included an automated XY stage controller and a humidified incubator set at 37 °C and 5% CO_2_.

### MTT assay

Cells cultured at 37 °C in 12-well plates were incubated in the dark for 75 min with culture medium containing 0.5 mg ml^-1^ MTT (Sigma-Aldrich). The cells were then carefully washed with PBS once and incubated for 10 min in the dark in dimethyl sulfoxide with mild shaking. Absorbance was measured at a wavelength of 570 nm using a microplate reader.

### Ex vivo 2-NBDG uptake assay

GSC–organotypic brain slice co-cultures were incubated overnight to ensure GSC engraftment into the brain slices. The co-culture brain slices were then incubated with 50 μM TAT or TAT-CX43_266-283_ for 48 h. On the day of the assay, the brain slices were incubated for 30 min in glucose-free medium (RPMI; Sigma-Aldrich) and subsequently for 1 h in glucose-free medium supplemented with 146 μM 2-NBDG. The brain slices were then washed with ice-cold PBS and mounted for confocal microscopy with SlowFade Gold Antifade reagent (Life Technologies). Images were taken on a Leica DM-IRE2 confocal microscope. For 6-NBDG uptake analysis, regions of interest (ROIs) were generated with the ‘Wand’ tool (ImageJ) in the GSC images and the mean grey values of the ROIs were measured in the 2-NBDG images.

### Intracranial implantation of glioma cells

Mice were anaesthetized by isoflurane inhalation, placed on a stereotaxic frame and window trephined in the parietal bone. A unilateral intracerebral injection in the right cortex was performed with a Hamilton microsyringe. Cellular suspensions were kept on ice while the surgery was being performed. Then, 1 μl of physiological saline containing 5,000 cells was injected into the cortex of adult mice placed in a stereotaxic frame at the following coordinates: 5 mm caudal to bregma, 4 mm lateral and 2 mm depth. Murine mCherry-G166 GSCs glioma cells or human G166 GSCs labelled with a Red Fluorescent Cell kit PKH26 (Sigma-Aldrich) were injected into NOD SCID mice. The needle was held in place for 1 min before being removed.

At the indicated times, the animals were transcardially perfused under deep anaesthesia (pentobarbital 120 mg kg^-1^, 0.2 ml) with 15 ml of physiological saline followed by 25 ml of 4% paraformaldehyde in 0.1 M phosphate buffer (pH 7.4). Brainstems were removed and cryoprotected by immersion in a solution of 30% sucrose in PBS until they sank. Then, 20-40-μm-thick coronal sections were obtained with a cryostat to be processed for immunostaining.

### Statistical analysis

The results are expressed as the means ± s.e.m. The number of technical replicates and independent experiments is indicated for each experiment in its corresponding figure or figure legend. For comparison between two groups, data were analysed by two-tailed Student’s t-test. When more than two groups were compared, data were analysed by one-way ANOVA, and confidence intervals (95%) and significance were corrected for multiple comparisons with the Tukey test. In all cases, values were considered significant when *P* < 0.05. Exact *P* values can be found in Supplementary Table 1.

## Acknowledgements

We thank Prof. J. Carretero (INCYL, University of Salamanca) for helping with the mouse surgery, T. del Rey and R. Flores (University of Salamanca) for the technical assistance, A. Álvarez Vázquez for helping with photomicrography and C. Ijurko, M. Romo-González and Á. Hernández (University of Salamanca) for helping with the qPCR experiments. This work was supported by the *Ministerio de Economía y Competitividad*, Spain; FEDER BFU2015-70040-R, FEDER RTI2018-099873-B-I00 and *Fundación Ramón Areces*. S.G. Pelaz and M. Jaraíz-Rodríguez are fellowship recipients from the *Junta de Castilla y León* and the European Social Fund. L. García-Vicente is a fellowship recipient from the *Ministerio de Ciencia, Innovación y Universidades* and R. Talaverón is a postdoctoral fellowship recipient from the University of Salamanca.

## Author contributions

S.G.P. contributed to the experimental design and development, data acquisition, analysis and interpretation of all of the experiments and drafted the article. S.G.P., M.J.-R., R.T. and L.G.-V. performed the in vivo experiments. M.G.C., M.T. and A.R.M. helped to design, acquire and process the data from the Seahorse experiments and supervised them. C.L. processed the samples for electron microscopy analysis and obtained and interpreted the images. J.M.M. contributed to the experimental design and data interpretation. A.T. conceived and designed the experiments, supervised the experimental development and analysis, interpreted the data and drafted the article. All of the co-authors revised the article for important intellectual content and approved the final version for publication.

