## Supplementary information for "TAT-Cx43_266-283_ impairs metabolic plasticity in glioma stem cells in vitro and in vivo"

### Supplementary Fig. 1

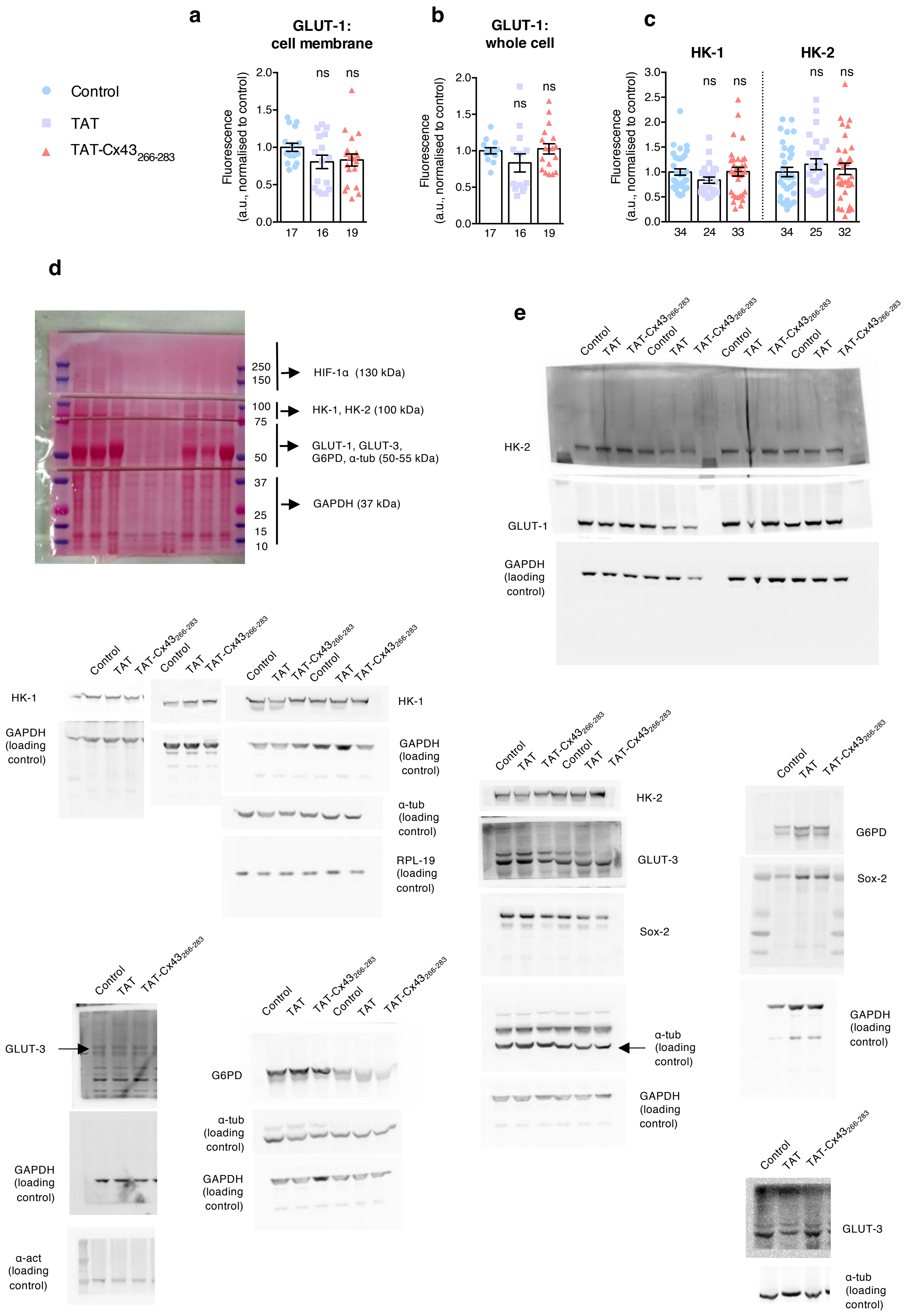

#### Supplementary Fig. 2

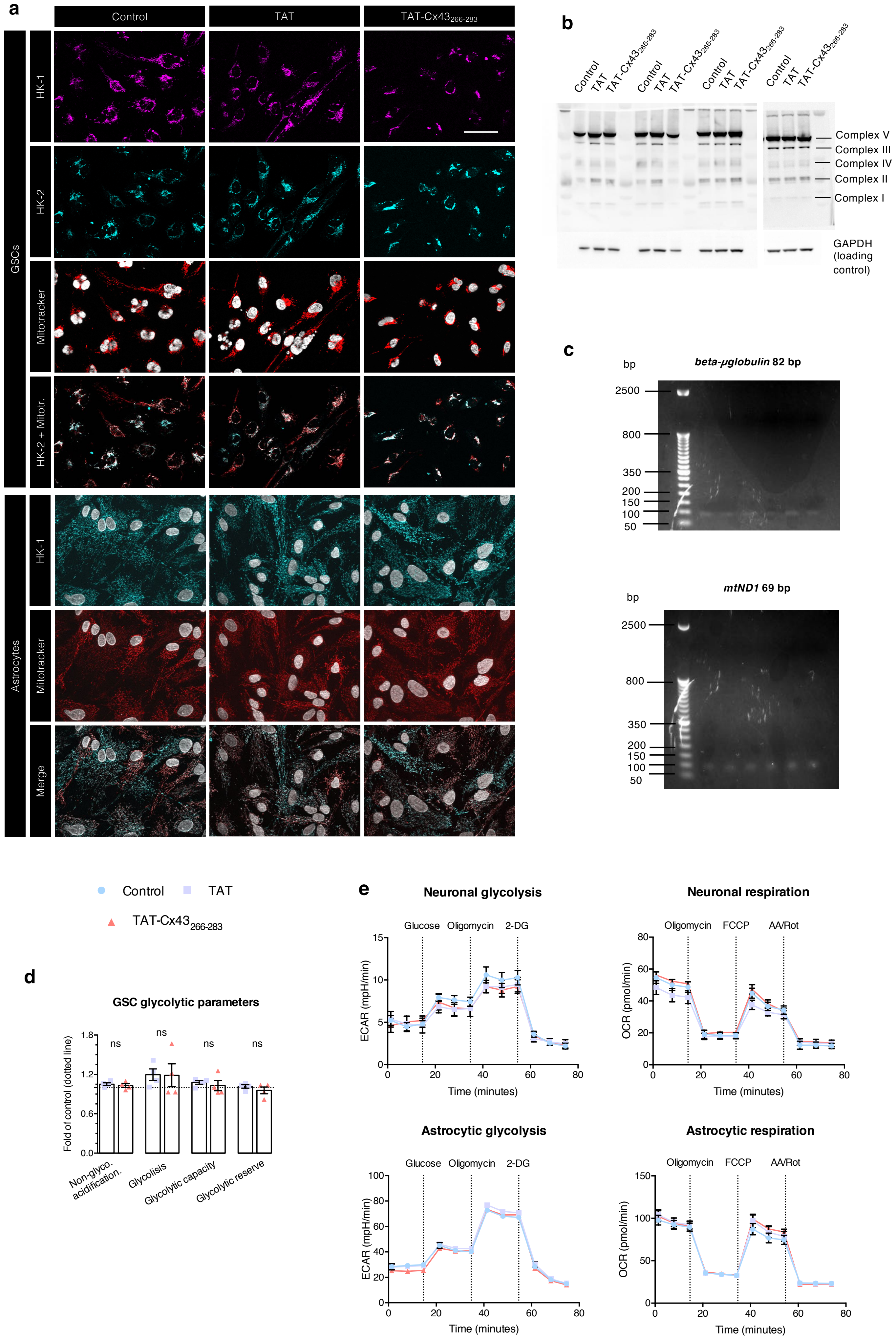

### Supplementary Fig. 3

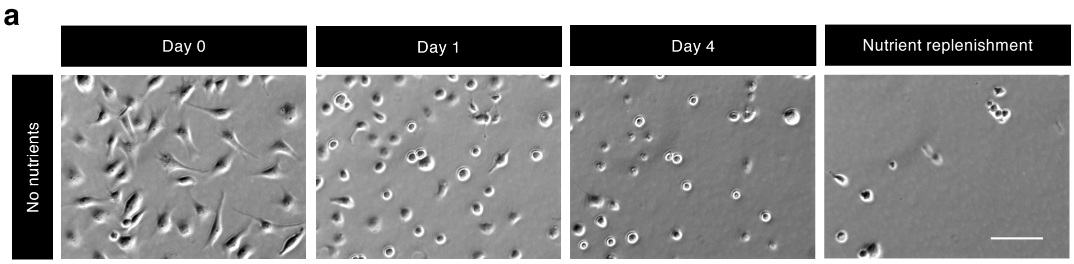


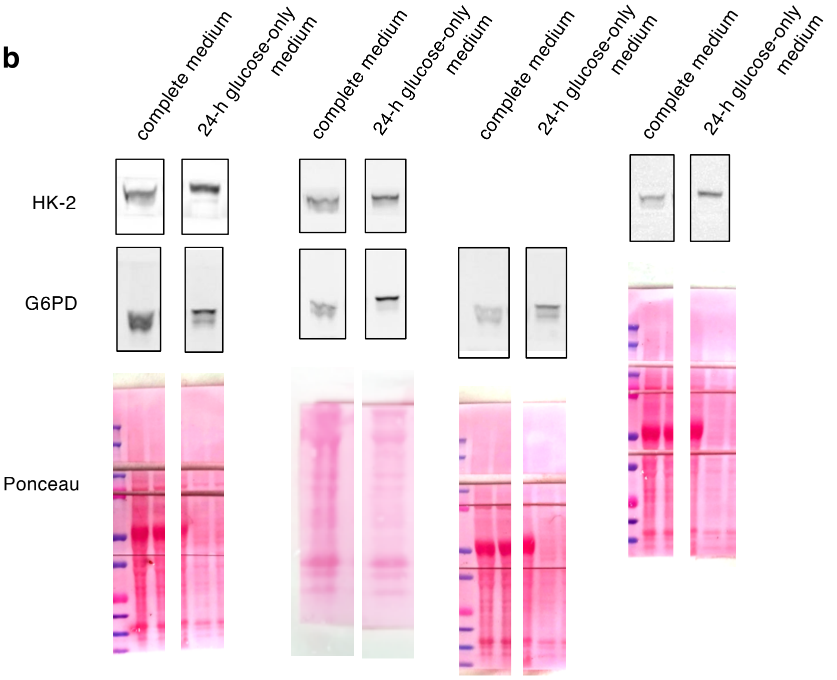


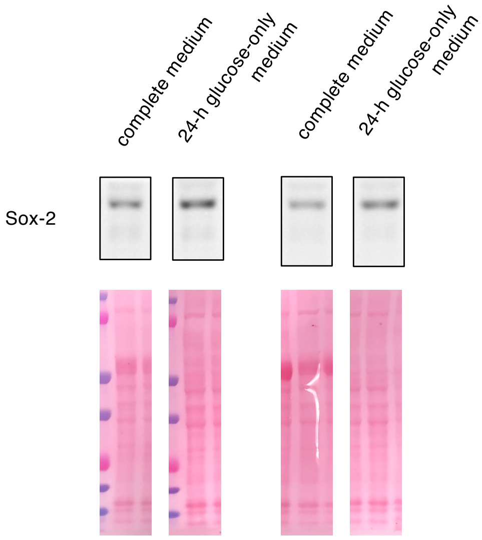

#### Supplementary Fig. 4

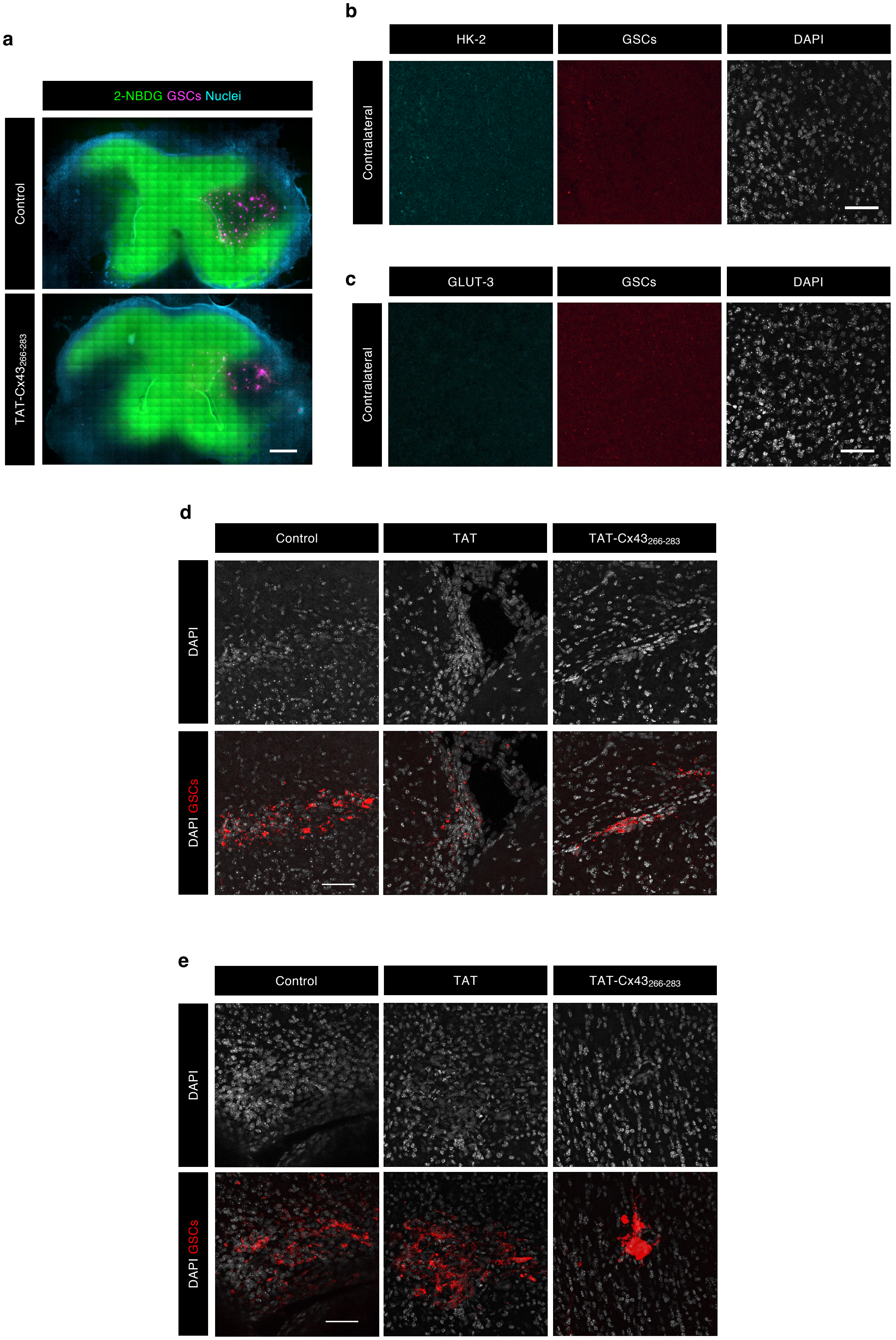

#### Supplementary Fig. 5

a

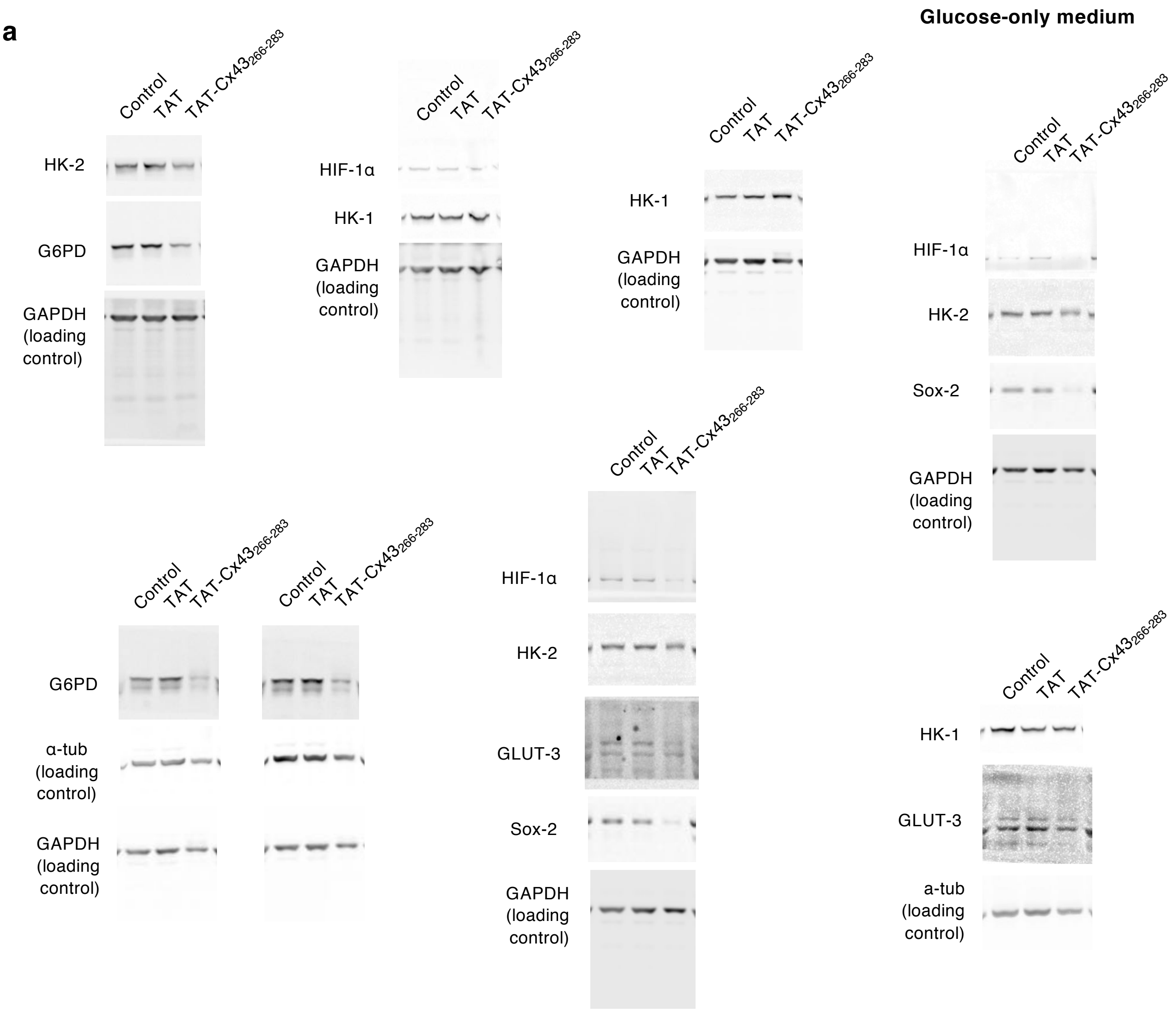


b

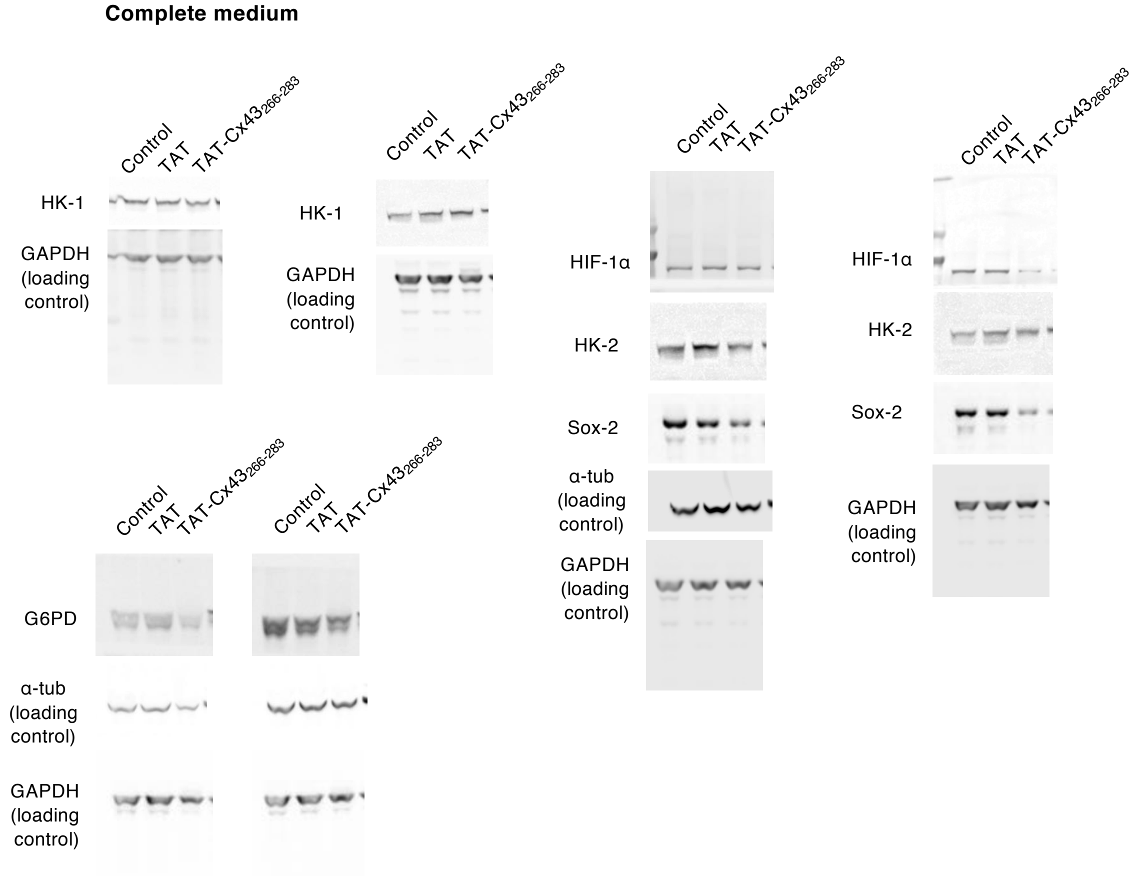

### Flg. 1-4

#### Fig. 1

| Fig. 1a | | Fig. 1d | | Fig. 1h | |
|---|---|---|---|---|---|
| **P value (ANOVA)** | 0,7222 | **P value (ANOVA)** | 0,0003 | **P value (ANOVA)** | 0,0119 |
| **C vs TAT** | 0,9969 | **C vs TAT** | 0,9839 | **C vs TAT** | 0,1759 |
| **C vs TAT-Cx43$_{266-283}$** | 0,7855 | **C vs TAT-Cx43$_{266-283}$** | 0,0016 | **C vs TAT-Cx43$_{266-283}$** | 0,0098 |
| **TAT vs TAT-Cx43$_{266-283}$** | 0,7435 | **TAT vs TAT-Cx43$_{266-283}$** | 0,0013 | **TAT vs TAT-Cx43$_{266-283}$** | 0,1124 |

| Fig. 1e - GSCs | | Fig. 1e - Neurons | | Fig. 1e - Astrocytes | |
|---|---|---|---|---|---|
| **P value (ANOVA)** | 0,0015 | **P value (ANOVA)** | 0,4225 | **P value (ANOVA)** | 0,9481 |
| **C vs TAT** | 0,8977 | **C vs TAT** | 0,5697 | **C vs TAT** | 0,9480 |
| **C vs TAT-Cx43$_{266-283}$** | 0,0036 | **C vs TAT-Cx43$_{266-283}$** | 0,9669 | **C vs TAT-Cx43$_{266-283}$** | 0,9685 |
| **TAT vs TAT-Cx43$_{266-283}$** | 0,0054 | **TAT vs TAT-Cx43$_{266-283}$** | 0,4307 | **TAT vs TAT-Cx43$_{266-283}$** | 0,9972 |

#### Fig. 2

| Fig. 2b | | Fig. 2c | |
|---|---|---|---|
| **P value (ANOVA)** | < 0.0001 | **P value (ANOVA)** | < 0.0001 |
| **C vs TAT** | 0,9997 | **C vs TAT** | 0,9649 |
| **C vs TAT-Cx43$_{266-283}$** | < 0.0001 | **C vs TAT-Cx43$_{266-283}$** | < 0.0001 |
| **TAT vs TAT-Cx43$_{266-283}$** | < 0.0001 | **TAT vs TAT-Cx43$_{266-283}$** | < 0.0001 |

| Fig. 2e | |
|---|---|
| **TAT vs TAT-Cx43$_{266-283}$ - P value (paired t-test, two-tails)** | 0,8001 |

| Fig. 2j | |
|---|---|
| **P value (paired t-test, two-tails)** | 0,2035 |

| Fig. 2h | Non-mito. resp. | Basal respiration | Proton leak | ATP-linked resp. | Maximal resp. capacity | Spare resp. capacity |
|---|---|---|---|---|---|---|
| **P value (ANOVA)** | 0,0790 | 0,0048 | 0,0064 | 0,0131 | 0,0304 | 0,0154 |
| **C vs TAT** | 0,1479 | 0,1510 | 0,0072 | 0,5056 | 0,9235 | 0,4764 |
| **C vs TAT-Cx43$_{266-283}$** | 0,9084 | 0,0418 | 0,7178 | 0,0488 | 0,0586 | 0,0139 |
| **TAT vs TAT-Cx43$_{266-283}$** | 0,0865 | 0,0040 | 0,0169 | 0,0125 | 0,0367 | 0,0412 |

#### Fig. 3

| Fig. 3b | Complete medium | Glucose-only medium | Amino acid-only medium | Glucose and amino acid medium | TAT-Cx43$_{266-283}$ in different media |
|---|---|---|---|---|---|

**Supplementary Figure Legends**

**Supplementary Fig. 1**. Related to Fig. 1.

(**a**) Quantification of GLUT-1 fluorescence intensity in the cell membrane. Images obtained with Apotome (optical sectioning structured illumination microscopy) were thresholded to exclude non-cell areas and the mean grey value of the resulting area was measured. Results were normalized to the control condition. (**b**) Quantification of GLUT-1 fluorescence intensity. Epifluorescence images from the same fields as (**a**) were acquired, thresholded to exclude non-cell areas and the mean grey value of the resulting area was measured. Results were normalized to the control condition. (**c**) Quantification of HK-1 and HK-2 fluorescence intensity. ROIs containing HK-1 or HK-2 signal were selected and measured. The values shown are the mean grey value of each ROI multiplied by its area (integrated density). Results were normalized to the control condition. (**d**) Blots were routinely cut into smaller pieces to evaluate different proteins from the same samples. Typically, blots were cut as shown in the Ponceau-stained image. (**e**) Original western blots and replicates from Fig. 1e. GAPDH, α-tubulin, α-actinin and RPL-19 blots are shown as loading controls. All data are mean ± s.e.m. and were obtained from at least three independent experiments with at least two technical replicates (ns, not significant). Numbers under bars indicate the number of images analysed in (**a**, **b**, **c**).

**Supplementary Fig. 2** Related to Fig. 2.

(**a**) Images showing HK-1 (magenta), HK-2 (cyan) and mitochondrial staining (MitoTracker, red) in GSCs; and HK-1 (cyan) and mitochondrial staining (MitoTracker, red) in astrocytes. Note that HKs and mitochondria have a similar distribution in all the cells and that, in GSCs treated with TAT-Cx43$_{266-283}$, HK-1, HK-2 and mitochondria are condensed near the cell nuclei, which does not occur in astrocytes. Scale bar: 50 µm. (**b**) Original western blots and replicates from Fig. 2d. GAPDH blots are shown as loading controls. (**c**) PCR products were run on a 2% agarose gel to ascertain correct primer amplification. (**d**) Quantification of glycolytic parameters calculated from data shown in Fig. 2g. Data from four independent experiments (ns, not significant). (**e**) Bioenergetic profiles of neurons and astrocytes from rat primary cultures. The OCR of neurons and astrocytes was measured after addition of 1.5 µM oligomycin to block ATP-linked OCR, 0.4 µM FCCP to uncouple mitochondria for the maximal OCR and 0.5 µM rotenone/antimycin A (Rot/AA) to shut down mitochondrial respiration. The ECAR was measured after addition of 10 mM glucose to assess the glycolysis rate, 1.5 µM oligomycin to obtain the maximal ECAR and 50 mM 2-DG to shut down glycolysis. Data from at least three independent experiments.

**Supplementary Fig. 3**. Related to Fig. 3.

(**a**) GSCs were recorded by time-lapse microscopy (Video 3) in medium without any nutrients. After 4 days, the medium was changed to complete medium and the cells were recorded for another 24 h (nutrient replenishment). Note that very few cells remain alive even after nutrient replenishment. Scale bar: 100 µm. (**b**) Western blots showing upregulation of G6PD, HK-2 and Sox-2 in GSCs cultured in glucose-only medium for 24 h. The same blots stained with Ponceau dye show the amount of protein loaded into each lane. Note the change in G6PD staining between the complete and glucose-only media, which suggests differential post-translational modifications of this protein that might affect its activity.

**Supplementary Fig. 4**. Related to Fig. 4.

(**a**) Representative mosaic photomicrographs of GSC–brain slice co-cultures incubated with 146 µM 2-NBDG. Note the similar levels of 2-NBDG uptake (green) by the brain parenchyma in control and TAT-Cx43$_{266-283}$–treated co-cultures. Scale bar: 1 mm. (**b** and **c**) Contralateral staining of GLUT-3 and HK-2 in brain sections of a xenograft mouse model of glioma. Note the absence of GSC staining (red) and human HK-2 (turquoise, **b**) or GLUT-3 (turquoise, **c**) staining in the contralateral hemispheres of the brain slices shown in Figure 5. Scale bar: 50 µm. (**d** and **e**) DAPI staining of the same fields shown in Fig. 4e (**d**) and Fig. 4g (**e**). Scale bar: 50 µm.

**Supplementary Fig. 5**. Uncropped Western blots from Fig. 3. GAPDH and α-tubulin blots are shown as loading controls.

(**a**) Western blots of GSCs treated with 50 µM TAT or TAT-Cx43$_{266-283}$ for 24 h in complete medium and then for 24 h in glucose-only peptide-containing medium showing a reduction in the levels of G6PD, GLUT-3, HIF-1α, HK-2 and Sox-2. HK-1 levels were not affected by the treatment. (**b**) Western blots of GSCs treated with 50 µM TAT or TAT-Cx43$_{266-283}$ for 24 h in complete medium and then for another 24 h in complete peptide-containing medium showing a mild reduction in the levels of G6PD. HK-1 levels were not affected by the treatment. Each blot is an independent experiment.

**Supplementary Fig. 6**

Names of all statistical analyses performed and the *P* values obtained.

#### Videos

**Videos 1–3.** GSCs were grown in glucose-only medium (Video 1.1), amino acid-only medium (Video 2.1) or without nutrients (Video 3.1) for 4 days. Frame rate= 25 frames per second. Then, the media were replaced by complete medium for 24 h (Videos 1.2, 2.2 and 3.2, respectively). Frame rate= 20 frames per second.

**Videos 4–9.** GSCs were treated with 50 µM TAT or TAT-$Cx43_{266-283}$ for 24 h and then the media was replaced with the indicated ones and then recorded for 24h. Videos 4 (complete medium) and 5 (glucose-only medium) correspond to the control condition, Videos 6 (complete medium) and 7 (glucose-only medium) correspond to the TAT condition and Videos 8 (complete medium) and 9 (glucose-only medium) correspond to the TAT-$Cx43_{266-283}$ condition. Frame rate= 5 frames per second.

| | | | | |
|---|---|---|---|---|
| **P value (ANOVA)** | < 0.0001 | < 0.0001 | 0,0331 | < 0.0001 |
| **C vs TAT** | 0,1908 | 0,7020 | 0,9924 | 0,1080 |
| **C vs TAT-$Cx43_{266-283}$** | < 0.0001 | < 0.0001 | 0,0573 | < 0.0001 |
| **TAT vs TAT-$Cx43_{266-283}$** | < 0.0001 | < 0.0001 | 0,0477 | < 0.0001 |

| | |
|---|---|
| **P value (ANOVA)** | < 0.0001 |
| **Complete vs glucose-only medium** | < 0.0001 |
| **Complete vs amino acid-only medium** | < 0.0001 |
| **Complete vs glucose and amino acid medium** | 0,0069 |
| **Glucose-only vs glucose and amino acid medium** | 0,0491 |
| **Amino acid-only vs glucose and amino acid medium** | 0,0211 |

| **Fig. 4** | |
|---|---|
| **Fig. 4b** | |
| **P value (ANOVA)** | < 0.0001 |
| **C vs TAT** | 0,8651 |
| **C vs TAT-$Cx43_{266-283}$** | < 0.0001 |
| **TAT vs TAT-$Cx43_{266-283}$** | < 0.0001 |

### Supplementary Fig. 1-5

#### Supplementary Fig. 1

**Fig. 1a**

| | |
|---|---|
| **P value (ANOVA)** | 0,1607 |
| **C vs TAT** | 0,1922 |
| **C vs TAT-$Cx43_{266-283}$** | 0,2530 |
| **TAT vs TAT-$Cx43_{266-283}$** | 0,9708 |

**Fig. 1b**

| | |
|---|---|
| **P value (ANOVA)** | 0,2340 |
| **C vs TAT** | 0,3898 |
| **C vs TAT-$Cx43_{266-283}$** | 0,9713 |
| **TAT vs TAT-$Cx43_{266-283}$** | 0,2337 |

**Fig. 1c**

| | HK-1 | HK-2 |
|---|---|---|
| **P value (ANOVA)** | 0,2414 | 0,5989 |
| **C vs TAT** | 0,3016 | 0,5692 |
| **C vs TAT-$Cx43_{266-283}$** | 0,9979 | 0,8983 |
| **TAT vs TAT-$Cx43_{266-283}$** | 0,2798 | 0,8232 |

#### Supplementary Fig. 2

**Supplementary Fig. 2c**

| | Non-glyco. acidification | Glycolysis | Glycolytic Capacity | Glycolytic Reserve |
|---|---|---|---|---|
| **P value (ANOVA)** | 0,2565 | 0,4214 | 0,5268 | 0,4623 |
| **C vs TAT** | 0,2311 | 0,4723 | 0,5031 | 0,9380 |
| **C vs TAT-$Cx43_{266-283}$** | 0,5904 | 0,4982 | 0,9014 | 0,6462 |
| **TAT vs TAT-$Cx43_{266-283}$** | 0,7303 | 0,9987 | 0,7552 | 0,4532 |

#### Supplementary Fig. 4

**Fig. 4c**

| | |
|---|---|
| **P value (ANOVA)** | 0,0178 |
| **C vs TAT** | 0,7903 |
| **C vs TAT-$Cx43_{266-283}$** | 0,0197 |
| **TAT vs TAT-$Cx43_{266-283}$** | 0,0431 |

**Fig. 4e**

| | |
|---|---|
| **P value (ANOVA)** | 0,0033 |
| **C vs TAT** | 0,7076 |
| **C vs TAT-$Cx43_{266-283}$** | 0,0039 |
| **TAT vs TAT-$Cx43_{266-283}$** | 0,0086 |
